# Gestational stress decreases postpartum mitochondrial respiration in the prefrontal cortex of female rats

**DOI:** 10.1101/2022.12.16.520624

**Authors:** Erin Gorman-Sandler, Breanna Robertson, Jesseca Crawford, Olufunke O. Arishe, R. Clinton Webb, Fiona Hollis

**Affiliations:** Department of Pharmacology, Physiology, and Neuroscience, University of South Carolina School of Medicine, Columbia, SC, USA; Columbia VA Health Care Systems, Columbia, SC, 29208, USA; Department of Cell Biology and Anatomy, University of South Carolina School of Medicine, Columbia, SC, USA; Cardiovascular Translational Research Center, University of South Carolina School of Medicine, Columbia, SC, USA

**Keywords:** chronic unpredictable stress, prefrontal cortex, pregnancy, postpartum, susceptibility, mitochondria

## Abstract

Postpartum depression (PPD) is a major psychiatric complication of childbirth, affecting up to 20% of mothers, yet remains understudied. Mitochondria, dynamic organelles crucial for cell homeostasis and energy production, share links with many of the proposed mechanisms underlying PPD pathology. Brain mitochondrial function is affected by stress, a major risk factor for development of PPD, and is linked to anxiety-like and social behaviors. Considering the importance of mitochondria in regulating brain function and behavior, we hypothesized that mitochondrial dysfunction is associated with behavioral alterations in a chronic stress-induced rat model of PPD. Using a validated and translationally relevant chronic mild unpredictable stress paradigm during late gestation, we induced PPD-relevant behaviors in adult postpartum Wistar rats. In the mid-postpartum, we measured mitochondrial function in the prefrontal cortex (PFC) and nucleus accumbens (NAc) using high-resolution respirometry. We then measured protein expression of mitochondrial complex proteins and 4-hydroxynonenal (a marker of oxidative stress), and Th1/Th2 cytokine levels in PFC and plasma. We report novel findings that gestational stress decreased mitochondrial function in the PFC, but not the NAc of postpartum dams. However, in groups controlling for the effects of either stress or parity alone, no differences in mitochondrial respiration measured in either brain regions were observed compared to nulliparous controls. This decrease in PFC mitochondrial function in stressed dams was accompanied by negative behavioral consequences in the postpartum, complex-I specific deficits in protein expression, and increased Tumor Necrosis Factor alpha cytokine levels in plasma and PFC. Overall, we report an association between PFC mitochondrial respiration, PPD-relevant behaviors, and inflammation following gestational stress, highlighting a potential role for mitochondrial function in postpartum health.

## 1. Introduction

Pregnancy and the postpartum period are accompanied by a high risk for complications which may negatively affect both mother and child. Mood alterations are a common complication, with depressive symptoms reported in 70% of women during pregnancy (Becker et al., 2016; O’Hara and Wisner, 2014). Furthermore, up to 1 in 5 women experience severe depressive symptoms within the year following parturition, often resulting in the diagnosis of postpartum depression (PPD) (“Depression Among Women | Depression | Reproductive Health | CDC,” 2020; O’hara and Swain, 1996). While PPD overlaps major depressive symptoms such as anhedonia, loss of energy, sleep disturbances, social dysfunction, cognitive impairment, feelings of hopelessness, and comorbid anxiety, it is further characterized by hallmark disturbances in the mother-infant relationship which include a lack of interest in the child, difficulty bonding, and in severe cases, thoughts of harm towards the self or child (O’Hara, 2009). As such, PPD accounts for the majority of postpartum deaths in the form of suicide (Becker et al., 2016) and is associated with impairments in maternal care which can negatively affect child development and increase their risk for psychiatric disturbances later in life (O’Hara, 2009). Although PPD contributes to increased health risks in both mother and child, with potentially long-term consequences, the underlying neurobiology of PPD remains unclear and effective treatment options are limited.

Stress is one of the largest risk factors for developing PPD, outside of a previous history of depression (Qiu et al., 2020), with negative, stressful life events associated with increased depressive symptoms (Lancaster et al., 2010). Moreso, women with PPD may have altered hypothalamic-pituitary-adrenal (HPA) axis function (Jolley et al., 2007). As such, animal models based on gestational chronic stress paradigms or corticosterone (CORT) injections administered during gestation or postpartum period induce PPD-relevant behaviors, such as altered maternal care and anhedonia (Brummelte and Galea, 2010; Leuner et al., 2014a; O’Mahony et al., 2006; Pardon et al., 2000; Perani and Slattery, 2014). These behavioral outputs have been associated with molecular changes in inflammatory responses, hormone signaling, neuroplasticity, and neurotransmission (Brummelte and Galea, 2016; Corwin et al., 2015; McEwen et al., 2012; Pawluski et al., 2016, 2011; Schiller et al., 2015). For example, stress-sensitive regions such as the prefrontal cortex (PFC), nucleus accumbens (NAc), and hippocampus (HPC) demonstrate structural changes and increased cytokine levels in animal models of postpartum depression (Dye et al., 2022; Haim et al., 2014; Leuner et al., 2014a; Workman et al., 2013). Additionally, the steroid hormone progesterone and its metabolite and neurosteroid, allopregnanolone, are associated with adaptation to stress (Brunton et al., 2014). Both women with PPD (Walton and Maguire, 2019) and rodents exposed to chronic stress (Bali and Jaggi, 2014) exhibit decreased levels of allopregnanolone. Recent work found that treatment with a synthetic formulation of allopregnanolone that targets GABAA receptors, Brexanolone, was highly successful in reversing PPD symptoms in humans (Kanes et al., 2017), though the long-term mechanism is unclear (Walton and Maguire, 2019) and the current treatment regimen is expensive and intensive. GABA hypofunction has been observed in both humans and rodent models of PPD (Maguire and Mody, 2008; Walton and Maguire, 2019) and treatment with Brexanolone acutely reverses this deficit in rodents (Melón et al., 2018). Interestingly, these molecular processes are all intimately linked to mitochondrial function (Accardi et al., 2014; Bordt et al., 2019; Bordt and Polster, 2014; Cheng et al., 2010; Du et al., 2009; West, 2017), suggesting a potential role for mitochondria in the stress-induced alterations observed in PPD.

Mitochondria are essential organelles that produce energy to facilitate cellular, physiological, and behavioral responses (Picard et al., 2018). The brain consumes approximately 20% of the body’s total energy (Rettberg et al., 2014), and stress can amplify that need, leading to increased mitochondrial activity to meet energy demands (Morava and Kozicz, 2013; Picard et al., 2018). Thus, mitochondria have been recently identified as vital for stress adaptation and may be involved in depressive pathology, with even small deficits in function producing impairments in synaptic plasticity and neurogenesis (Brinton, 2008a; Morava and Kozicz, 2013; Picard et al., 2018; Rettberg et al., 2014). Preclinical studies in male rodents found that chronic stress reduced mitochondrial function in the cortex and cerebellum (Madrigal et al., 2001; Rezin et al., 2008). Importantly, however, the effects of gestational stress and parity on brain mitochondria in postpartum females remains completely unexplored. We previously demonstrated a direct role for mitochondrial function within the NAc and PFC in anxiety-like and social behaviors (Hollis et al., 2018, 2015) revealing an active role for these organelles in regulating behavioral phenotypes relevant to depression. Moreover, mitochondria release immunogenic compounds that stimulate the release of pro-inflammatory cytokines (West and Shadel, 2017) that have been implicated in PPD (Dye et al., 2021). Combined with the involvement of mitochondria in many of the physiological changes that occur during pregnancy and parturition, as well as their vital role in stress and inflammation, mitochondria have great potential to mediate postpartum-induced alterations in brain and behavior, and may contribute to susceptibility in the development of PPD.

We sought to determine whether gestational stress exposure impacts mitochondrial function in specific brain regions implicated in PPD and mitochondrial mediation of behavior. We exposed pregnant female rats to chronic mild unpredictable stress (CMUS) during late gestation, and assessed postpartum behaviors relevant to PPD, brain mitochondrial respiration, and cytokine levels. We hypothesized that CMUS will induce postpartum depressive-like behaviors and decrease brain mitochondrial function within stress-sensitive regions implicated in PPD, specifically the PFC and NAc that may associate with increased cytokine levels. We found that gestational CMUS induced a PPD-relevant phenotype, and interestingly, decreased mitochondrial respiration in the PFC but not in the NAc. These findings provide rationale to consider mitochondrion as a therapeutic target for postpartum depression.

## 2. Methods

### 2.1 Animals

Intact and cycling nulliparous and timed mated primiparous adult female Wistar rats (Envigo, Virginia, USA) initially weighing 225-250g arrived at the vivarium in the same delivery day and were used in these experiments. Thus, experimental time points for both nulliparous and timed-mated females are matched to the primiparous female gestational day (GD). Upon arrival on gestational day 4 (GD4), rats were housed individually in standard polycarbonate rat cages on a 12:12h light/dark cycle (lights on at 07:00h) and provided food and water ad libitum. Single-housing allowed for individual home cage assessments of behavior (such as the sucrose preference test and maternal care). All animals were weighed on arrival and handled for at least three days before the start of any manipulations. Stress manipulations began 1 week after arrival to allow for habituation to the vivarium (see Figure 1 for an experimental timeline).

**Figure 1.**
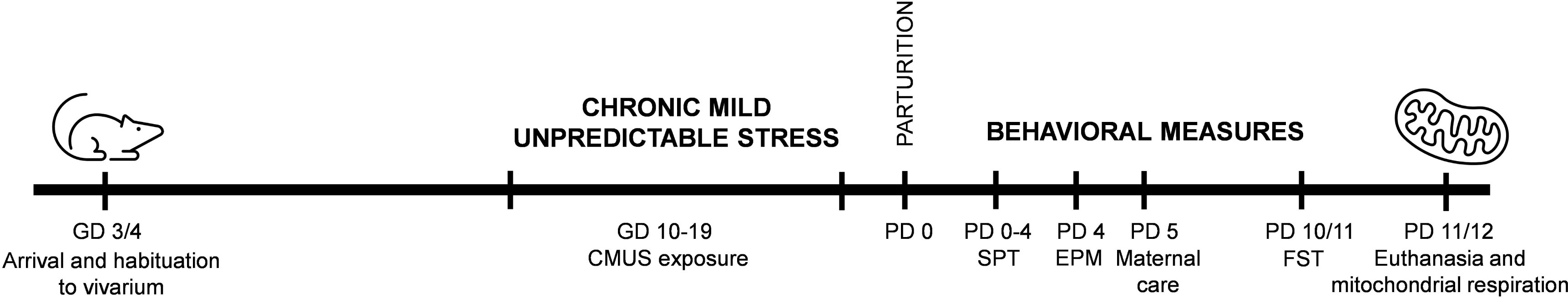
Experimental Timeline. Nulliparous and primiparous female rats were weighed and handled upon arrival at the vivarium. Females were weight-matched (within parity groups) and assigned to stress or non-stress conditions. Chronic mild unpredictable stress (CMUS) began on the day corresponding to gestational day (GD) 10 in primiparous rats and continued for 10 days through GD 19. Following parturition on GD 21/postpartum day (PD) 0, a series of behavioral tests were performed during the postpartum period. The sucrose preference test (SPT) was performed from PD 0-4, the elevated plus maze (EPM) was performed on PD 4, maternal care was assessed during the onset of the dark cycle on PD 5, and the one-session forced swim test (FST) was performed on PD 10 or 11. All rats that underwent behavior were euthanized on the days corresponding to PD 11 or 12 and mitochondrial respiration was measured in the prefrontal cortex and nucleus accumbens. Other molecular analyses (protein and cytokine measurements) are representative of this same euthanasia timepoint.

Animals were weight-matched and assigned to the following experimental groups:

1. Nulliparous/non-stressed (control: C)
2. Nulliparous/stressed (stress only: S)
3. Primiparous/non-stressed (primiparous only: P)
4. Primiparous+stressed (P+S)

The inclusion of C, S, and P groups allow us to account for potential individual effects of stress and pregnancy on weight, behavior, and mitochondrial respiration. Groups exposed to stress were housed in a separate room from non-stressed groups. After the start of stress, all animals were weighed every other day and cage changes were performed twice weekly, except on the days surrounding behavioral testing. During behavioral testing, cage changes were avoided the day before a behavioral measure to limit potential effects on behavior. All behavioral measurements were performed between 08:00 and 14:00h, with the exception of the home cage sucrose preference test and maternal care observations (see description below under “Sucrose Preference Test” and “Maternal Care”). The day of parturition was designated as postpartum day 0 (PD 0). Litters were characterized briefly for number of pups and sex ratio (Table 1). All efforts were made to minimize animal suffering and reduce the number of animals used where possible. All experimental procedures were approved by The University of South Carolina Institutional Animal Care and Use Committee and conformed to the U.S. National Institutes Health Guide for the Care and Use of Laboratory Animals.

**Table 1.** Litter characteristics. Primiparous dams had no significant differences in litter size, litter weight, sex ratio, or mortality rate (*Two-tailed unpaired Student’s t-test*; *Welch’s correction* applied for analysis of pup mortality).

| Measure | P (n=9) | Std dev. | P+S (n=9) | Std dev. | p-value | t-value | df |
| --- | --- | --- | --- | --- | --- | --- | --- |
| Avg litter size (#) | 11.2 | 4.969 | 11.7 | 3.606 | 0.8308 | 0.2172 | 16 |
| Avg litter weight (g) | 8.6 | 0.805 | 8.4 | 0.6815 | 0.5437 | 0.618 | 20 |
| Avg percentage of male offspring (%) | 46.5 | 11.72 | 55.03 | 16.6 | 0.2359 | 1.235 | 15 |
| Pup mortality (#) | 0 | 0 | 0.1111 | 0.3333 | 0.3466 | 1 | 8 |

### 2.2 Chronic Mild Unpredictable Stress (CMUS)

Nulliparous (S) and primiparous (P+S) groups were exposed to a chronic, mild, unpredictable stress (CMUS) protocol, adapted from Frisbee *et al*. This protocol has emerged as a highly translationally-relevant model for studying the pathophysiology of chronic stress exposure in rodents (Willner, 2017; Willner et al., 1987) A fundamental concept of the CMUS protocol that enhances its relevance to human situations lies in the heterotypic, unpredictable and uncontrollable nature of the stressors that prevents adaptation and leads to disrupted stress response systems (Frisbee et al., 2015). Our protocol consisted of a randomized series of daily exposures to one of the following stressors (Table 2): white noise, bedding alterations (wet bedding, no bedding), tilted cage, overnight light exposure, and acute restraint stress during the second half of gestation (GD 10-GD 19), to avoid altered reproductive effects (Gemmel et al., 2018). The length of exposure for each stressor, as well as the time of day that each was introduced was varied to maximize unpredictability and uncontrollability. Both S and P+S groups began stress exposure on the same day and experienced the same order and timing of stressors. The last stress manipulation occurred on GD 19. PPD-relevant behaviors were measured following parturition, after cessation of stress.

**Table 2.** Chronic Mild Unpredictable Stress paradigm. Nulliparous and primiparous female rats in the stress group began stress exposure on the day corresponding to GD 10. Stress exposure occurred over varying lengths of time and at different times of day, and included wet bedding, no bedding, overnight light, white noise, cage tilt, and restraint stress, as detailed in the table. All cohorts followed the same stress schedule.

| Day | Stressor | Duration |
| --- | --- | --- |
| GD10 | Wet bedding | 3h |
| GD11 | Tilted cage (30° angle) | 5h |
| GD12 | Overnight light | 12h |
| GD13 | Restraint stress | 3h |
| GD14 | Bedding removed | 4h |
| GD15 | Wet bedding | 7h |
| GD16 | White noise | 6h |
| GD17 | Overnight light | 12h |
| GD18 | Restraint stress | 1h |
| GD19 | Wet bedding | 4h |

### 2.3 Behavioral Testing

#### 2.3.1 Sucrose Preference Test

The sucrose preference test, a two-bottle choice paradigm, was performed according to the procedure outlined by Bolaños *et al*. (Bolaños et al., 2008) and our lab (Hollis et al., 2011, 2010). This test has been used extensively in evaluating stress-induced anhedonia (Willner, 2017; Willner et al., 1987), a core symptom of depression characterized by lack of ability to experience pleasure. Rats were habituated to drink water from two 250 mL bottles for four days. A water intake baseline was measured for 48 hours within the week prior to the start of sucrose introduction in order to assess differences in fluid intake. From PD0-4, all groups were exposed to ascending concentrations of sucrose (0.25% and 1%) for two days per concentration. Animals were group matched, and the exact timing of when the test began was dependent on time of parturition. Notably, all pregnant dams gave birth within 24 hours of each other. The amount of water and sucrose solution consumed was measured daily by weighing bottles. The positions of the bottles in each cage were switched between the groups 1-3 times daily to eliminate effects of side preference. The preference for sucrose over water was used as a measure of the rats’ response to a naturally rewarding stimulus. Specifically, sucrose preference was measured as (sucrose intake (grams)/total water + sucrose intake (grams)) x 100%. Preference significantly lower than controls was considered anhedonic behavior.

#### 2.3.2 Elevated Plus Maze

On PD 4, all groups were tested for anxiety-like behavior in the elevated plus maze. Rats were habituated to the testing room for at least 30 minutes. The apparatus was elevated (50 cm) from the floor and consisted of two closed arms (50 × 10 × 40 cm), two open arms (50 × 10 cm), and was separated by a center zone (10 × 10 cm). Lighting was maintained at 4-7 lux in the closed arms, 18-21 lux on the open, and 12-15 lux in the center. These lighting conditions ensure observations of trait anxiety as opposed to state anxiety, as very bright lighting can itself be anxiogenic (Hollis et al., 2015). Animals were placed in the center of the maze facing a closed arm, and behavior on the maze was recorded for 300s. Before testing and in between trials, the apparatus was cleaned with a 10 % EtOH solution. Distance moved, average velocity, and time spent in the center, open arms, and closed arms were measured. Anxiety-like behavior was determined by % time spent in the open arms. (Time spent in open arms (sec)/total time of test(sec)) x 100%. Behaviors were scored using automated scoring by EthoVision XT.

#### 2.3.3 Maternal Care: Dark-cycle home cage nesting observations

Home cage nesting observations were used to assess maternal behaviors in basal conditions on PD 5, as alterations in maternal care are characteristic of a gestationally stressed phenotype. Dams were recorded for one hour at the start of the dark cycle (19:00-20:00h) using a video recorder under red light conditions. Cages of P and P+S dams were turned with the nest facing the video recorder. The experimenter was not present during the recording period, and videos were scored at a later time. Scored behaviors included number of on-nest and off-nest events. Active on-nest events included licking/grooming of pups, and nursing. Scoring was performed by an experimenter blind to the experimental conditions using Noldus Observer XT15. Videos were instantaneously sampled every minute across the 60-minute session and 60 total events (behaviors) were recorded and analyzed.

#### 2.3.4 Forced Swim Test

The forced swim test was performed on the day preceding euthanasia (PD10 or 11) to characterize passive versus active coping style. Active coping is associated with more time spent swimming/struggling, while passive is associated with more time spent immobile (Molendijk and de Kloet, 2015). Animals were brought to the behavioral room to habituate for at least 30 minutes. Cylindrical tanks (60 cm in height) were filled with fresh water, at a temperature between 24-26°C. Animals were individually placed into the tank and underwent swim exposure for a total of 15 minutes. After the 15-minute test, animals were taken out of the tank and dried by towel and space heater before being returned to their individual home cages. The entire procedure was video-recorded, and behavior was later quantified manually by an experimenter blind to the treatments, for latency to become immobile and time spent immobile. Latency to immobility was defined as the time at which the rat first adopted a stationary pose not associated with an attempt to escape. Immobility was scored when the rat remained in this stationary posture for more than 2.0 s, making only the movements necessary to maintain its head above water.

### 2.4 Euthanasia and tissue collection

Animals were euthanized on PD 11 or12 under basal conditions by rapid decapitation. Importantly, this mid-postpartum time point corresponds with a return of estradiol levels to basal levels, following a peak at the end of gestation that steadily decreases through PD 14 (Camacho-Arroyo et al., 2018). Trunk blood was collected in EDTA-coated tubes (Sarstedt) and immediately chilled in ice. Plasma was isolated from blood via centrifugation at 16,000xg for 3 min. Plasma was immediately snap frozen in chilled isopentane and stored at -80°C until further processing. The brain was rapidly removed and PFC and NAc dissected out. One hemisphere from each region was snap frozen in chilled isopentane and stored at -80°C. The remaining hemisphere was weighed and placed in a well plate on ice with 2 mL of relaxing solution (BIOPS: 2.8 mM Ca2K2EGTA, 7.2 MK2EGTA, 5.8 mM ATP, 6.6 mM MgCl2, 20 mM taurine, 15 mM sodium phosphocreatine, 20 mM imidazole, 0.5 mM dithiothreitol and 50 mM MES, pH = 7.1) until further processing.

### 2.5 Mitochondrial Respirometry

Tissue samples were processed as described in (Hollis et al., 2015). Briefly, samples were gently homogenized in ice cold MiR05 respirometry medium with a motorized Teflon pestle. One hemisphere (approximately 20 mg) of tissue was used to measure mitochondrial respiration rates at 37°C using high resolution respirometry (Oroboros Oxygraph 2K, Oroboros Instruments, Innsbruck, Austria). A multi-substrate protocol was used to sequentially explore the various components of mitochondrial respiratory capacity as previously described (Hollis et al., 2015). The O_2_ flux obtained in each step of the protocol was normalized by the wet weight of the tissue sample used for the analysis and corrected for ROX activity. In line with other published reports (Burtscher et al., 2015), we use CICII/ETS-linked capacity as an internal standardization factor to control for differences in mitochondrial density between samples.

### 2.6 Protein expression

Relative protein expression was assessed using Western blotting technique, with primary antibodies (AB) against individual subunits of each complex in the oxidative phosphorylation (OXPHOS) system, 4-hydroxynonenal (4-HNE), a marker of oxidative stress, and Translocase of the Outer Membrane 20 (TOM20), a receptor protein located on the outer mitochondrial membrane. Protein concentration of PFC homogenates in respirometry medium was determined by BCA assay (ThermoScientific, Cat. #23227). Samples were prepared with Laemelli buffer and milliQ water to a final concentration of 1µg/µL, then heated in a dry bath for 5 minutes at either 37°C (for OXPHOS AB detection) or 95°C (for 4-HNE and TOM20 AB detection). 20 µL of protein lysate was then loaded into 4–20% Criterion™ TGX Stain-Free™ precast gels (Bio-Rad, Cat. #5678094). Proteins were separated at 200V for 45 min and then transferred to a low-fluorescence PVDF membrane at 25V for 30 min using the Trans-Blot Turbo Transfer System and Kit (Bio-Rad, Cat. #1704275). Gels were activated by UV excitation using the Bio-Rad GelDoc XR+ System prior to transfer, and membranes were imaged for total protein on the same system following transfer. Membranes were then blocked with EveryBlot Blocking Buffer (Bio-Rad Cat. #12010020) for 30 min (for 4-HNE and TOM20 AB detection) or 1 hr (for OXPHOS AB detection) and then incubated overnight at 4°C with Rodent Total OXPHOS antibody cocktail (Mitosciences, ab110413; 1:250), anti-4-Hydroxynonenal (Sigma-Aldrich, AB5605; 1:4000), or TOM20 (Abclonal, A19403; 1:2000) in blocking buffer. Following washes in PBS-T (VWR, Cat. #76371-736), membranes were incubated with IRDye® 680RD Goat-anti-Mouse (LI-COR, #926-68070) or IRDye® 800CW Donkey anti-Goat (LI-COR, #926-32214), or IRDye® 800 CW Goat anti-Rabbit (LI-COR, #926-32211) IgG secondary antibodies at a 1:20000 dilution for 1h at room temperature. After PBS-T washes, bands were captured using a LI-COR Odyssey CLX Imager. Bands were quantified as 16-bit TIF images using ImageJ software (NIH). The 4-HNE antibody binds to HNE-modified protein adducts resulting from lipid peroxidation. Thus, this antibody results in multiple bands within each lane, which were all quantified. Quantified bands (whole lane quantified for 4-HNE) were normalized to total protein to control for variance in sample loading. Western blot data are presented as fold change relative to control averages to control for between-blot variability.

### 2.7 Assessment of central and peripheral cytokine levels

Cytokine levels were assessed using Bio-Rad BioPlex assays in both plasma and PFC homogenates (Th1/Th2 rat cytokines Bio-Rad, #171k1002M), according to manufacturer’s protocols. PFC homogenates were diluted 1:3 and plasma were diluted 1:4 with diluent. Plates were read on a Luminex plate reader using high photomultiplier voltage and analyzed with Bio-Plex manager software. PFC cytokine levels were normalized to tissue protein.

### 2.8 Statistical Analyses

Sample sizes are indicated in the figure legends. Data were analyzed by unpaired Student’s t-test (maternal care and litter parameters) or two-way ANOVAs (all other behaviors, protein expression data, and cytokine data), followed by *Šídák’s* or Dunnett’s multiple comparisons post hoc test where appropriate. Two-way ANOVAs tested stress exposure and pregnancy as between subject factors. Correlational data were analyzed using a simple linear regression. Figures are presented with original, non-transformed data. All data were analyzed using Prism version 9 (GraphPad software Inc., San Diego, CA). P-values are reported in figure legends. Statistical significance was considered at the p<0.05 level.

## 3. Results

### 3.1 Validation of gestational CMUS paradigm

We first validated that our CMUS protocol successfully induced PPD-relevant behaviors. In rodent studies, several groups report that exposure to gestational stress induces reduced gestational weight gain, anhedonia, reduced or disrupted maternal care, and passive coping in the forced swim test (Haim et al., 2014; O’Mahony et al., 2006; Pardon et al., 2000; Smith et al., 2004; Zhang et al., 2022; Zoubovsky et al., 2020). Some report evidence of increased anxiety-like behavior (Darnaudéry et al., 2004; Hillerer et al., 2011), while others observe no effect (Leuner et al., 2014b). Thus, we selected these same behaviors for assessment in our animals to ensure that our application of the CMUS protocol was effective.

We recorded weight gain across CMUS exposure, observing an expected significant main effect of parity on weight gain during the stress period, with primiparous groups gaining more total weight than nulliparous groups due to gestation (Fig 2A). Within parity groups, primiparous+stress (P+S) dams had significantly reduced total body weight gain compared to their non-stressed primiparous (P) counterparts, while nulliparous control (C) and stress (S) groups did not differ from each other (Fig 2A). Despite differences in weight gain, there were no differences in litter size, sex ratio, pup survival rate, or average pup weight (at PND 4) between primiparous dams (Table 1).

**Figure 2.**
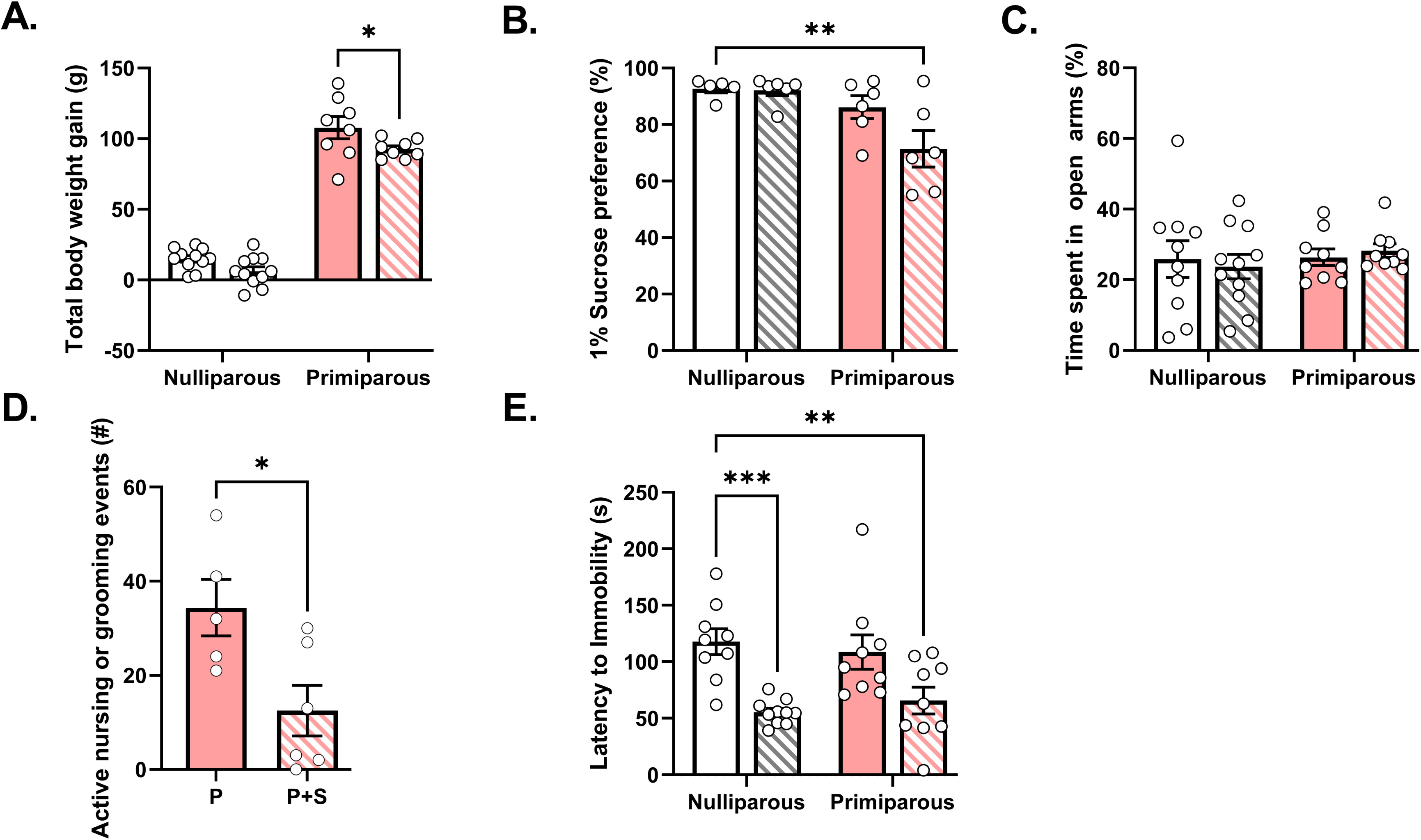
CMUS successfully induced PPD-relevant behaviors in gestationally stressed dams. **(A)** P+S dams had significantly lower body weight gain during GD 10-19 (stress period) than P dams, while S females did not significantly differ in body weight gain from C females, as evidenced by significant main effects of both stress and parity (*two-way ANOVA*; stress: F_(1,34)_ = 8.489; p = 0.0063; parity: F_(1,34)_ = 476.5; p<0.0001), but no significant interaction between stress and parity (F_(1,34)_ = 0.5890; p = 0.4481). (*Šídák’s multiple comparisons test;* P vs. P+S: p = 0.0417, C vs. S: p = 0.2033). n = 8-11/group. **(B)** There was a significant main effect of parity on sucrose preference (*two-way ANOVA;* F_(1,19)_= 10.72; p = 0.004), and trends for a main effect of stress (*two-way ANOVA;* F_(1,19)_ = 3.368; p = 0.0822), where P+S dams exhibited significantly lower preference for a 1% sucrose solution compared to nulliparous controls (*Dunnett’s multiple comparisons test*; C vs. P+S: p = 0.006). Neither P nor S groups demonstrated decreased sucrose preference compared to C rats (*post-hoc*; C vs. P: p = 0.5624, C vs. S: p = 0.9993). There was no statistically significant interaction between stress and parity in 1% sucrose preference (F_(1,19)_ = 2.882; p = 0.1059). n = 5-6/group. **(C)** There was no significant interaction between stress and parity on percent time spent in open arms, and no main effects of either parity or stress (*two-way ANOVA;* interaction: F_(1,35)_ = 0.3053; p = 0.5841, parity: F_(1,35)_ = 0.4763; p = 0.4946, stress: F_(1,35)_ = 0.0006; p = 0.9814). n = 9-11/group. **(D)** P+S dams had significantly reduced number of active on-nest events compared to P dams (*Two-tailed unpaired Student’s t-test;* t = 2.7171, df = 9; p = 0.0237). n = 5-6/group. **(E)** There was a significant main effect of stress on both nulliparous and primiparous females in latency to immobility in the forced swim test (*two-way ANOVA;* F_(1,33)_ = 27.17; p < 0.0001), but no significant interaction or main effect of parity (interaction: F_(1,33)_ = 1.33; p = 0.2571, parity: F_(1,33)_ = 0.0324; p = 0.8583). Both S and P+S groups had significantly shorter latencies to become immobile compared to C rats, while P dams showed no significant difference in latency to immobility compared to C rats (*Dunnett’s multiple comparisons test*; C vs. S: p = 0.0002, C vs. P: p = 0.6748, C vs. P+S: p = 0.0018). n = 9-10/group. Nulliparous controls =C, nulliparous stressed =S, primiparous controls =P, and primiparous stressed =P+S. All data are represented as mean ± SEM. *p≤0.05, **p≤0.01, ***p≤0.001, ****p≤0.0001.

Following parturition, we assessed anhedonia in the early postpartum period using the sucrose preference test. Baseline measurement of water consumption revealed a significant effect of parity with both P and P+S females consuming more water than nulliparous groups (Supp Fig 1A). Assessment with 0.25% sucrose revealed trends for a significant effect of parity and interaction with stress (Supp Fig 1B), however, preference was low across all groups and not significantly different from 50%. When we measured preference for a higher concentration of sucrose solution (1%), we observed preferences significantly above the 50% indifference point (Fig 2B). We also observed a significant main effect of parity and trends for an effect of stress and interaction. Post-hoc analyses revealed that these effects were primarily driven by the P+S group, which exhibited significantly lower preference for a 1% sucrose solution compared to nulliparous controls (Fig. 2B). Neither P nor S rats demonstrated differences in sucrose preference compared to nulliparous controls (Fig 2B). Furthermore, there were no significant differences in total liquid consumption between groups (Supp Fig. 1C), ruling out differences due to overall consummatory behavior. We then assessed early postpartum anxiety-like behavior in the elevated plus maze. We found no significant interaction between stress and parity on percent time spent in open arms, and no main effects of either parity or stress. (Fig. 2C). Additionally, there were no significant interactions or main effects of stress or parity on the number of entries into open arms, distance traveled, or average velocity (Supp Fig 2A-C). Home cage maternal care observations during the active period revealed that P+S females had significantly fewer bouts of nursing and grooming compared to P counterparts (Fig 2D). We also evaluated stress-coping strategy in a one-day, 15-minute forced swim test in the mid-postpartum. We found a significant main effect of stress on both nulliparous and primiparous females in latency to immobility in the forced swim test, such that both S and P+S females had shorter latencies to immobility compared to nulliparous controls (Fig. 2E). There was no main effect of parity nor interaction. Non-stressed primiparous dams showed no significant difference in latency to immobility compared to nulliparous controls. Taken together, these data replicate previously published findings and demonstrate that our CMUS protocol was effective in inducing PPD-relevant behaviors.

### 3.2 Gestational stress reduces mitochondrial respiration in the prefrontal cortex that correlates to passive coping behavior

We next evaluated the effects of stress and parity on mitochondrial function in the postpartum prefrontal cortex (PFC) and nucleus accumbens (NAc), two brain regions implicated in PPD (Pawluski et al., 2017; Ray et al., 2016). Using high-resolution respirometry, we found that gestational stress induced significant reductions in PFC mitochondrial respiration. Specifically, gestational stress decreased coupled respiration through complex I (CI) and complex II (CI+CII; Fig. 3A). When respiration was uncoupled, gestational stress significantly reduced the maximal electron transport system capacity (Max; Fig. 3A). Upon complex I inhibition by rotenone, there were trends for maximal respiration due to complex II activity (Max + CII) to differ under stress and parity, suggesting that effects are driven by both complex I and II-dependent deficits in respiration (Fig. 3B). We analyzed CI&II-linked maximum capacity, an indication of mitochondrial content (Burtscher et al., 2015), and found no effects of parity or stress on mitochondrial content (Fig. 3C), suggesting that the observed respiratory chain deficits were not due to differences in mitochondrial number. Similarly, analysis of mitochondrial outer membrane protein TOM20 further revealed no differences between groups (Supp Fig 3A). Interestingly, we found that the latency to immobility in primiparous females was predictive of PFC complex I+II coupled respiration (Fig. 3D), suggesting a relationship between passive coping and PFC mitochondrial respiration. We did not observe any significant effects of parity or stress on mitochondrial respiration in the NAc across any of our measures (Fig. 3E and Supp Fig 3B), suggesting that the effects of gestational stress on mitochondrial function are not global.

**Figure 3.**
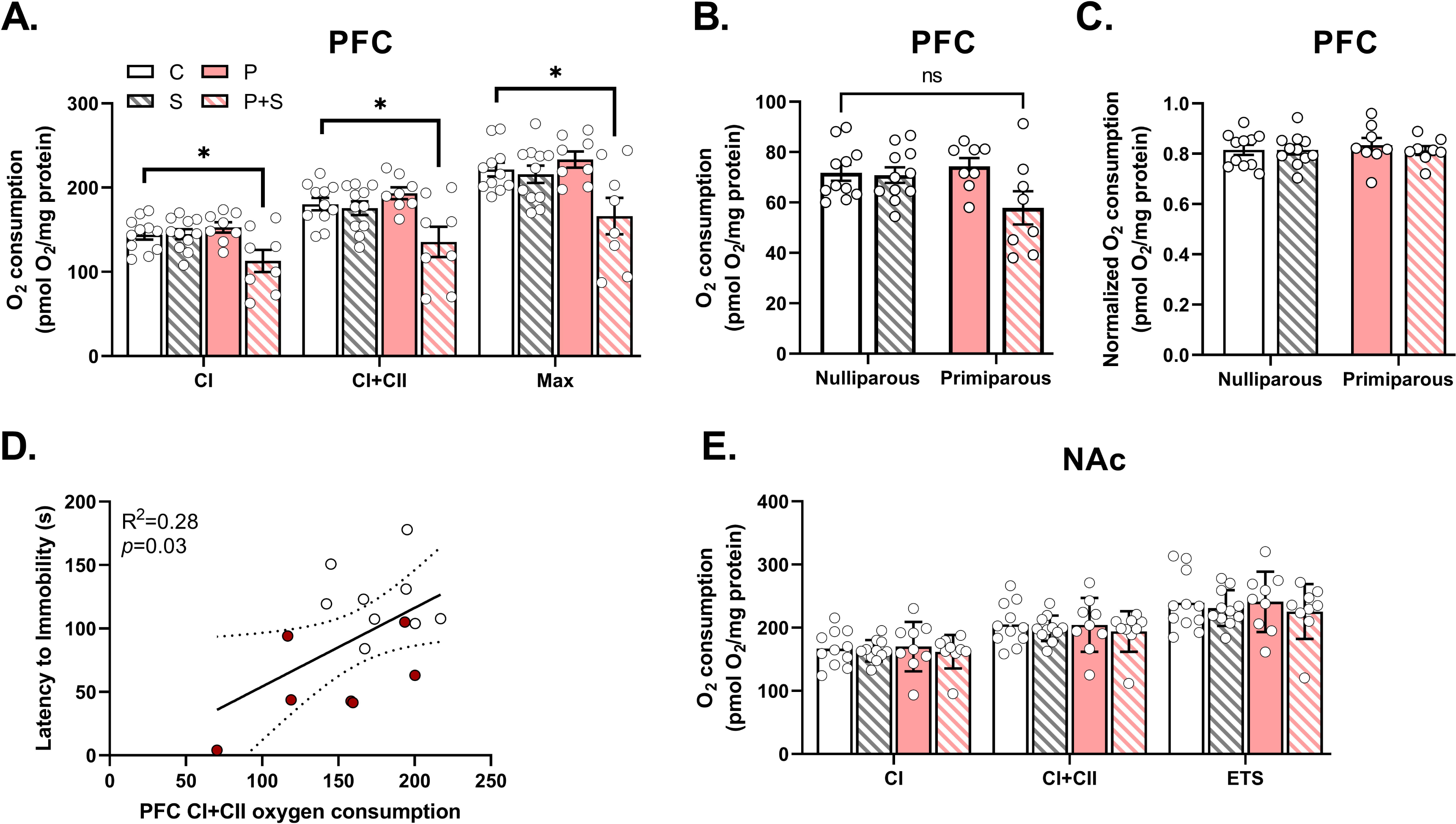
Gestational stress decreases mitochondrial respiration in the prefrontal cortex but not nucleus accumbens. **(A)** In prefrontal cortex (PFC), there was a significant interaction and main effect of stress, with no main effect of parity, on **complex I (CI)** (*two-way ANOVA;* interaction: F_(1,34)_ = 6.649; p = 0.0144, stress: F_(1,34)_ = 6.255, p = 0.0174, parity: F_(1,34)_ = 2.178, p = 0.1492), **complex I+II combined (CI+CII) (**interaction: F_(1,34)_ = 6.503; p = 0.0154, stress: F_(1,34)_ = 9.168; p = 0.0047, parity: F_(1,34)_ = 1.682; p = 0.2034) , and **maximal (MAX)** (interaction: F_(1,34)_ = 5.899; p = 0.0206, stress: F_(1,34)_ = 8.323; p = 0.0067, parity: F_(1,34)_ = 2.237; p = 0.1439) respiration, where post-hoc analyses demonstrated significantly decreased respiration in all measures in P+S dams compared to C females (*Dunnett’s multiple comparison’s test;* **(CI)** C vs. S: p = 0.9999, C vs. P: p = 0.7862, C vs. P+S: p = 0.0223. **(CI+CII)** C vs. S: p = 0.9693, C vs. P: p = 0.7179, C vs. P+S: p = 0.121. (**MAX)** C vs. S: p = 0.9730, C vs. P: p = 0.8555, C vs. P+S: p = 0.0109). n = 8-11/group. **(B)** In prefrontal cortex, there was a trend for an interaction between stress and parity on **complex II** respiration, with a significant main effect of stress and no main effect of parity (*two-way ANOVA;* interaction: F_(1,34)_ = 3.708; p = 0.0625, stress: F_(1,34)_ = 4.672, p = 0.0378, parity: 1.64; p = 0.209), although post-hoc analyses reveal a trend for a difference between P+S and C females (*Dunnett’s multiple comparison’s test* C vs. S: p = 0.9960, C vs. P: p = 0.9443, C vs. P+S: p = 0.0541). n = 8-11/group. **(C)** There were no significant differences observed between groups in mitochondrial content, as measured by ETS-linked/CI+CII respiration (*two-way ANOVA;* interaction: F_(1,34)_ = 0.2416; p=0.6262, parity: F_(1,34)_ = 0.1271 p= 0.7237, stress: F_(1,34)_ = 0.2336, p = 0.6319). n = 8-11/group. **(D)** Latency to immobility in P+S dams and C females was found to be predictive of complex I+II coupled respiration in the prefrontal cortex (*Simple linear regression*, R^2^ = 0.2773, p = 0.0361, where quicker latency to become immobile was associated with decreased oxygen consumption. Red dots = P+S dams, white dots = C females. n = 16. **(E)** In nucleus accumbens (NAc), there were no significant interactions or main effects of stress or parity on **complex I (CI) (***two-way ANOVA;* interaction: F_(1,36)_ = 0.0673; p = 0.7968, parity: F_(1,36)_ = 0.0067; p = 0.9353, stress: F_(1,36)_ 0.414, p = 0.5240), **complex I+II (CI+CII) (**interaction: F_(1,36)_ = 0.0453; p = 0.8327, parity: F_(1,36)_ = 0.0706; p = 0.7919, stress: F_(1,36)_ 0.6349, p = 0.4308), or **maximal (MAX)** (interaction: F_(1,36)_ = 0.0742; p = 0.7869, parity: F_(1,36)_ = 0.0206; p = 0.8866, stress: F_(1,36)_ = 0.8036; p = 0.3760) respiration. n = 9-11/group. Nulliparous controls =C, nulliparous stressed =S, primiparous controls =P, and primiparous stressed =P+S. All data are represented as mean ± SEM. *p≤0.05, **p≤0.01, ***p≤0.001, ****p≤0.0001.

### 3.3 Gestational stress blocks the parity-induced increase in mitochondrial complex I protein expression, with no effect on ROS

Given the reduction in mitochondrial respiration in the PFC, we sought to determine whether gestational stress decreased mitochondrial complex protein levels. Immunoblots of PFC homogenates revealed a significant interaction between stress and parity on complex I levels, such that primiparous non-stressed females tended to have higher complex I expression compared to nulliparous controls (Fig. 4A-B). Interestingly, P+S females exhibited significantly reduced complex I protein levels compared to P counterparts, suggesting that gestational stress prevented the increase in complex I protein following parturition. We did not observe any significant effects of parity, stress, or interaction on complex II, III, IV, or V protein levels (Fig. 4C-F), suggesting that the effect of parity and stress on mitochondrial complex protein may be specific for complex I. As stress has been shown to increase reactive oxygen species (ROS) and complex I is both a major site of ROS production and target for detrimental ROS actions (Musatov and Robinson, 2012), we also examined the PFC for evidence of enhanced levels of 4-hydroxynonenal (4-HNE), a primary ROS product. We did not observe any significant effects of stress, parity, or interaction on 4-HNE levels (Fig. 4G-H), indicating that the observed decrease in mitochondrial respiration is not due to enhanced lipid peroxidation.

**Figure 4.**
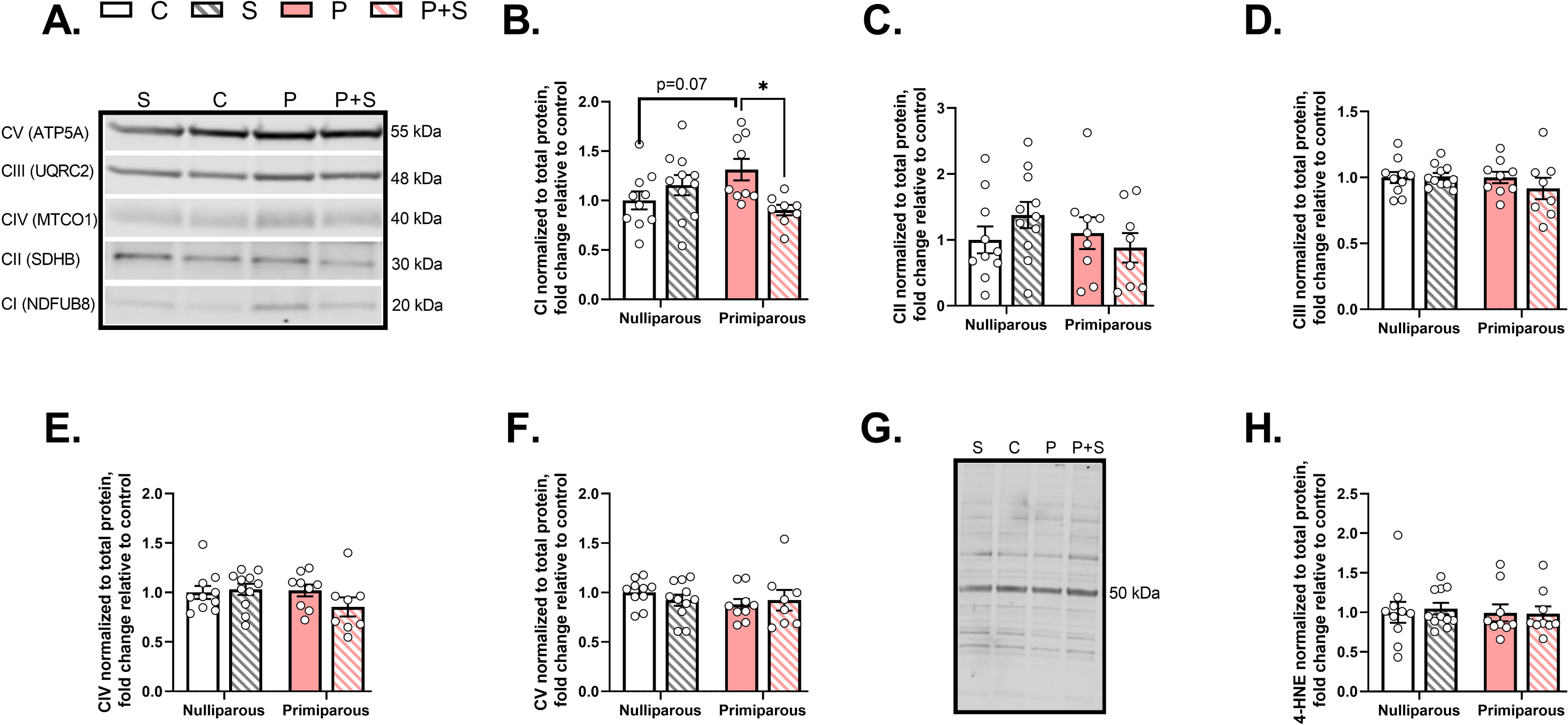
Gestational stress decreases mitochondrial complex I protein but has no effect on other complexes or 4-HNE protein expression in prefrontal cortex. **(A)** Representative Western blots of complex protein. **(B)** In prefrontal cortex, we found a significant interaction between stress and parity on complex I protein expression (*two-way ANOVA;* F_(1,34)_ = 8.741; p = 0.0056), with no significant main effects of parity (F_(1,34)_ = 0.0938, p = 0.7613) or stress (F_(1,34)_ = 1.777, p = 0.1914), such that P dams exhibited a trend for increased protein expression compared to C females (*Dunnett’s multiple comparison’s test* C vs. S: p = 0.4902, C vs. P: p = 0.0686, C vs. P+S: p = 0.8243), and P+S dams had significantly reduced protein expression compared to P dams (*Šídák’s multiple comparisons test* C vs. S: p = 0.4118, P vs. P+S: p = 0.0135). No significant differences were observed between groups in protein expression of **(C) complex II** (*two-way ANOVA*; interaction: F_(1,34)_ = 1.906; p = 0.1764, parity: F_(1,34)_ = 0.8231; p = 0.3707, stress: F_(1,34)_ = 0.1268; p = 0.7239), (D) complex III *two-way ANOVA*; interaction: F_(1,34)_ = 0.8882; p = 0.3526, parity: F_(1,34)_ = 0.8878; p = 0.3527, stress: F_(1,34)_ = 0.6035; p = 0.4426), **(E) complex IV** (*two-way ANOVA*; interaction: F_(1,34)_ = 2.05; p = 0.1613, parity: F_(1,34)_ = 1.266; p = 0.2685, stress: F_(1,34)_ = 0.9753; p = 0.3304), or **(F) complex V** (*two-way ANOVA*; interaction: F_(1,34)_ = 0.8478; p = 0.3637, parity: F_(1,34)_ = 0.8717; p = 0.3571, stress: F_(1,34)_ = 0.0691; p = 0.7943). Similarly, **(G)** representative western blot of 4-HNE revealed no significant effects of stress, parity, or interaction on **(H) 4-HNE** (*two-way ANOVA*; interaction: F_(1,35)_ = 0.0795; p = 0.7797, parity: F_(1,35)_ = 0.1181; p = 0.7332, stress: F_(1,35)_ = 0.0282; p = 0.8676). n = 8-11/group. Nulliparous controls =C, nulliparous stressed =S, primiparous controls =P, and primiparous stressed =P+S; 4-HNE= 4-hydroxynonenal. All data are represented as mean ± SEM. *p≤0.05, **p≤0.01, ***p≤0.001, ****p≤0.0001.

### 3.4 Gestational stress increases pro-inflammatory cytokine levels in the PFC and plasma that correlate with PFC mitochondrial respiration

PPD has been linked with inflammation (Anderson and Maes, 2013; Dye et al., 2022) and mitochondrial dysfunction can result in the release of immunogenic compounds that activate inflammasomes to promote pro-inflammatory cytokine release (West and Shadel, 2017). Therefore, we examined cytokine levels in the plasma and PFC homogenates using BioPlex assays. We found a significant main effect of stress on Tumor Necrosis Factor-alpha (TNF-α), with P+S specifically exhibiting significantly higher levels than C females (Fig. 5A). We did not observe any significant effects on levels of Interleukin (IL) 1-β, 1-α, 6, or 10 (Supp Fig. 4A-D). As these measures were taken from the same animals where mitochondrial respiration was measured, we performed Pearson correlations between mitochondrial respiration and TNF-α levels. We found a significant negative correlation between PFC TNF-α levels and coupled CI+CII respiration in stressed rats, such that higher levels of TNF-α were associated with lower mitochondrial coupled respiration (Fig. 5B). In contrast, we observed a somewhat different cytokine profile in the plasma. In plasma, we found significant main effects of parity on levels of TNF-α (Fig. 5C), as well as levels of IL-1β, IL-6, IL-18, IFN-γ, and VEGF (Supp Fig. 5A-E). Given the increased TNF-α in both brain and periphery, we performed correlations to determine if plasma TNF-α levels might be associated with central TNF-α levels and/or PFC mitochondrial respiration. Surprisingly, we did not observe a significant correlation between plasma and PFC TNF-α levels (Supp Fig. 5F). However, we did observe a significant negative correlation between plasma TNF-α and PFC mitochondrial respiration in primiparous females (Fig. 5D), highlighting an exciting possible relationship between peripheral TNF-α levels and central mitochondrial function.

**Figure 5.**
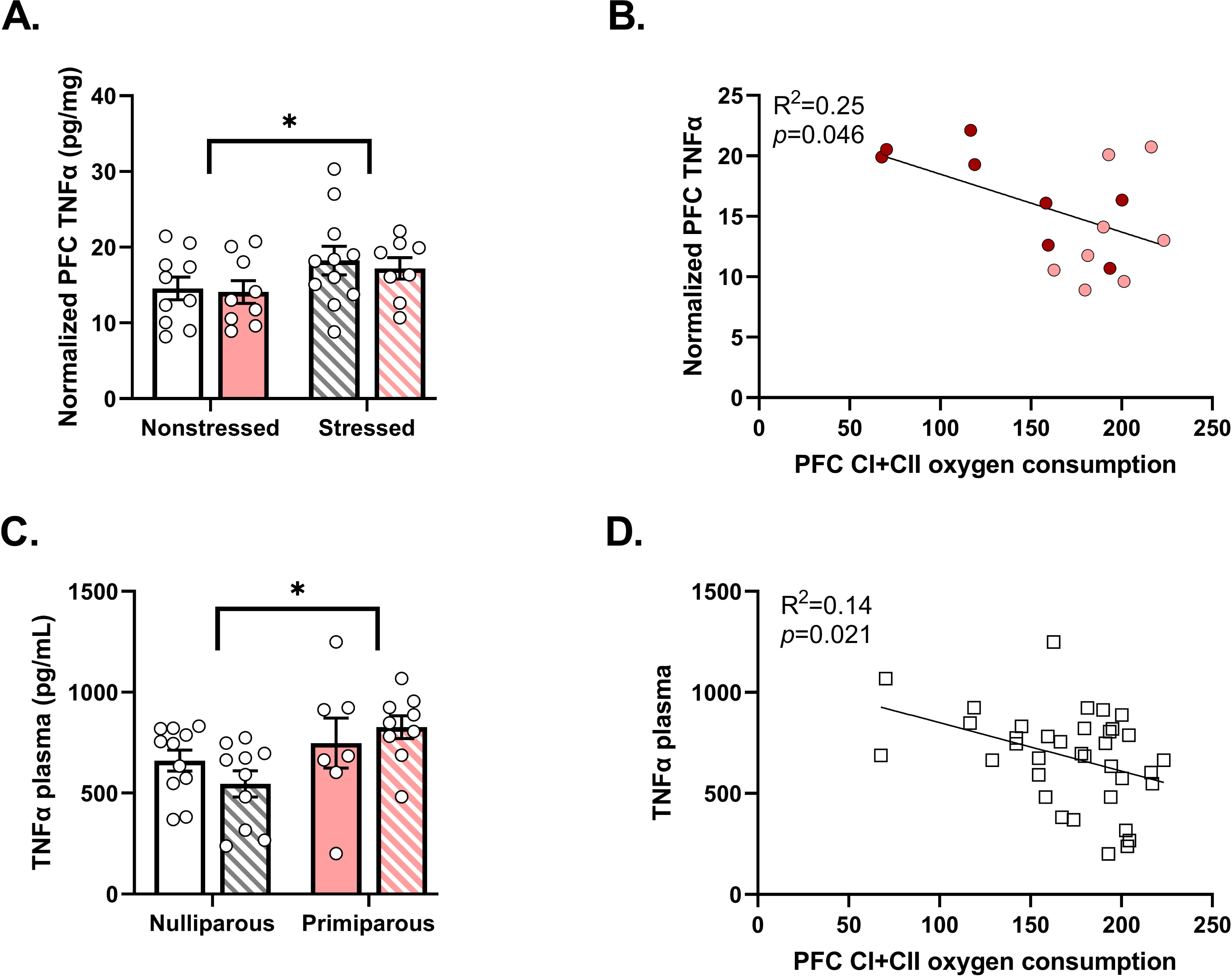
Gestational stress decreases tumor necrosis factor alpha (TNFα) levels in PFC and plasma. **(A)** We found a significant main effect of stress on TNFα in the prefrontal cortex, (*two-way ANOVA*; stress: F_(1,34)_ = 4.226; p = 0.0.0475). There was no significant main effect of parity (F_(1,34)_ = 0.2071; p = 0.652), nor interaction (*two-way ANOVA:* interaction: F_(1,34)_ = 0.0312; p = 0.8608). Post-hoc analyses demonstrated no significant differences in groups compared to C females (*Dunnett’s multiple comparisons test;* C vs. S: p = 0.2448, C vs. P: p = 0.9946, C vs. P+S: p = 0.564). n = 8-11/group. **(B)** In P (pink dots) and P+S (red dots) dams, CI+CII coupled respiration in the PFC was predictive of TNFα in the PFC, where higher levels of the cytokine correlated with lower oxygen consumption (*Simple linear regression;* n = 16.) **(C)** In plasma, there was a significant main effect of parity on TNFα, while there was no significant interaction or main effect of stress (*two-way ANOVA*; parity: F_(1,33)_ = 6.549; p = 0.0153, interaction: F_(1,33)_ = 1.813; p = 0.1873, stress: F_(1,33)_ = 0.0661; p = 0.7987). However, post-hoc analyses revealed no significant differences between groups (*Dunnett’s multiple comparisons test;* C vs. S: p = 0.5429, C vs. P: p = 0.7934, C vs. P+S: p = 0.2637). n = 7-11/group. **(D)** We observed that CI+CII coupled respiration in the prefrontal cortex was predictive also of TNFα levels in plasma, with inclusion of all groups, such that higher levels of plasma TNFα were associated with lower levels of oxygen consumption. (*Simple linear regression*. n = 36). Nulliparous controls =C, nulliparous stressed =S, primiparous controls =P, and primiparous stressed =P+S. All data are represented as mean ± SEM. *p≤0.05, **p≤0.01, ***p≤0.001, ****p≤0.0001.

## 4. Discussion

Here we show that increased postpartum depressive-like behaviors in dams exposed to chronic gestational stress is associated with decreased postpartum mitochondrial respiration in the prefrontal cortex (PFC) and increased inflammation at the central and systemic levels. In contrast, mitochondrial respiration in the nucleus accumbens (NAc) was unaffected by gestational stress exposure or pregnancy. Our inclusion of both nulliparous and primiparous stress and non-stressed groups allowed us to dissect the individual effects of stress and pregnancy as well as their interaction on behavior, postpartum brain mitochondrial function, and inflammation. While mitochondrial involvement has been shown in other behaviors, including social dominance (Hollis et al., 2015; van der Kooij et al., 2018), stress (Weger and Sandi, 2018), anxiety (Filiou et al., 2011; Filiou and Sandi, 2019), and depression (Allen et al., 2021), our study is the first to characterize brain mitochondrial functional changes both in the postpartum period and following exposure to chronic gestational stress. Overall, these data highlight the potential for mitochondria to act as a mediator for the pathological effects of stress on behavior and inflammation in the postpartum period.

First, using a late-gestational, 10-day chronic mild unpredictable stress paradigm, we induced maternal deficits, anhedonic, and passive stress-coping behaviors in female rats in the early postpartum period. Interestingly, we observed an interaction between stress and pregnancy in the sucrose preference – where only P+S females exhibited a decreased sucrose preference. While many studies using similar stress protocols have induced anhedonia in nulliparous animals (Franceschelli et al., 2014), in our hands CMUS exposure alone did not significantly affect sucrose preference. The most plausible explanation for this difference is the duration of our stress exposure, as a study by Baker *et al*. demonstrated limited effects of 3 weeks of chronic mild stress on nulliparous female sucrose preference, (Baker et al., 2006) and most CMUS studies range from 3 weeks to 3 months in duration, employing multiple stress exposures in a single day (Franceschelli et al., 2014). Our protocol duration was necessarily limited to 10 days to restrict exposure to the late gestational period, with a single stress exposure per day. We did not observe any effects of parity or stress on anxiety-like behavior in the Elevated Plus Maze, similar to some groups (Leuner et al., 2014b), but in contrast to others (Darnaudéry et al., 2004; Hillerer et al., 2011). A likely explanation for this discrepancy is a difference in test lighting conditions, as we measure in low light to assess trait rather than state anxiety and studies suggest the need for high lighting levels to evoke anxiety in nulliparous control females (Lonstein, 2007). It is important to note, however, that while symptoms of anxiety are often comorbid with those of depression in human populations, they certainly do not always occur together (Austin et al., 2010; Ross et al., 2003), and thus our CMUS protocol might serve to model PPD specifically outside of anxiety. Importantly, both nulliparous and primiparous stressed females exhibited significantly decreased latencies to immobility in the forced swim test, in line with other reports using rodent stress models, and suggesting that our stress protocol was successful. The finding that our stress paradigm elicited specific effects (*e.g.*, reduced weight gain and anhedonia) in primiparous, but not nulliparous, females suggests that pregnancy may increase susceptibility to developing maladaptive behaviors in this brief context of chronic mild stress.

The major finding of our study was the observation of specific differences in postpartum PFC mitochondrial function following gestational stress exposure that was associated with postpartum depression-relevant behaviors. Gestationally stressed (P+S) but not primiparous non-stressed (P) or nulliparous stressed (S) dams exhibited a significant decrease in postpartum mitochondrial respiration in the PFC. This decrease was not due to decreased mitochondrial number as mitochondrial content was similar between groups, as evidenced by a lack of difference between groups in PFC CI and CII-linked ETS capacity and TOM20 protein levels. Instead, we observed a significant decrease in complex I protein levels compared to P females. P females tended to exhibit increased postpartum complex I protein levels compared to nulliparous controls (C), while P+S females notably did not. The difference in protein levels is intriguing as P females did not exhibit any difference in respiratory levels from controls, highlighting the potential for a compensatory increase in protein expression to meet potentially increased metabolic demand in the postpartum period. While such studies in pregnant and postpartum women are limited, there is some evidence of pregnancy-driven physiological regulation of mitochondrial function, with pregnancy upregulating membrane potential to compensate for increased ROS production (Feldthusen et al., 2014). Notably, we did not observe any alterations in ROS lipid peroxidation products between any of our groups. This may be due to the analysis of different ROS products (4HNE vs TBARS, superoxides, or hydrogen peroxide production), or the specificity for cellular compartments. As we analyzed the total homogenate, it is possible that mitochondrial-specific effects were diluted. Taken together, our results suggest decreased PFC postpartum mitochondrial respiration in P+S females due to a lack of enrichment in complex I respiratory complex per mitochondria, compared to their P counterparts.

Interestingly, CMUS had no effect on mitochondrial respiration within the NAc of dams, indicating that postpartum brain mitochondrial respiration is not globally reduced by gestational stress. The NAc is largely involved in reward and motivation and plays a major role in maternal care during the postpartum, including pup-motivated behaviors (Champagne, 2004). Animal models of PPD typically demonstrate alterations in maternal care and have been linked to disruptions in NAc regulation (Champagne and Meaney, 2006a, 2006b; Pardon et al., 2000; Post and Leuner, 2019; Smith et al., 2004). Furthermore, in a study by Haim *et al*., chronic restraint stress between gestational days 7-20 induced changes in NAc structural plasticity during both the early/mid postpartum period (Haim et al., 2014). However, we did not observe any effects of parity or gestational stress on mitochondrial function within the NAc. It is possible that different types of stress can elicit varying phenotypes (Du Preez et al., 2021). For example, Haim and colleagues used homotypic chronic restraint stress while we employed a heterotypic stress protocol, using unpredictable stressors at different times of the day. It is also possible that a longer duration of stress would induce alterations that we do not observe with a 10-day protocol, or that differences would be observed in an earlier or later postpartum time point. Finally, there could be population-specific changes that we are unable to detect as we did not sub-divide the NAc into core and shell regions. Future studies that examine the effects of different types of stressors, additional time points, or specific cellular populations will allow us to further understand the specificity of effects of gestational stress on brain mitochondrial function.

The lack of effect in NAc also highlights the PFC as a region particularly susceptible to the remodeling effects of gestational CMUS. The PFC has consistently been implicated in preclinical studies in rodent models of stress and depression (Duman, 2014) and associated with decreased volume, neuronal atrophy, and altered connectivity in stress-related pathologies (Drevets et al., 2008). Furthermore, our findings corroborate other rodent studies reporting brain mitochondrial susceptibility to chronic stress. For instance, a study by Weger *et al*. observed chronic stress-induced mitochondrial gene signature alterations accompanied by reduced respiration in the PFC but not the NAc (Weger and Sandi, 2018). While these studies were primarily in males, used a longer duration of stress, and did not explore the contributions of pregnancy, they similarly support higher susceptibility of the PFC compared to the NAc to chronic stress, which could account for the decreased respiration we observe under our CMUS paradigm. Importantly, the PFC is one of several known brain regions affected by pregnancy, exhibiting volumetric (Hoekzema et al., 2017) and morphological (Leuner and Gould, 2010) changes suggestive of a role in maternal behaviors (Hillerer et al., 2014), and involvement in clinical (Moses-Kolko et al., 2010) and preclinical (Leuner et al., 2014b) models of PPD. Moreover, other studies using gestational stress models have reported enduring reductions in PFC dendritic spine density and morphology that was associated with PPD-relevant behaviors such as maternal care and passive coping (Haim et al., 2016; Leuner et al., 2014b). Mitochondria localize to synapses to provide crucial energy and buffering capacity during synaptic activity (Rangaraju et al., 2019) and artificial depletion of mitochondria results in decrease in dendritic spines and synapse density (Li et al., 2004). Thus, our observed reductions in mitochondrial respiration may underlie the stress-induced structural remodeling reported by others, though further studies are necessary to concretely link these two processes.

Immune system alterations have been noted in both pregnancy and the postpartum period. While normal pregnancy is characterized by an anti-inflammatory state with pro-inflammatory shifts depending on the stage (Bränn et al., 2019), stress during pregnancy can alter the immunological profile of mothers, promoting a more pro-inflammatory state that is associated with PPD (Anderson and Maes, 2013; Christian, 2015). Stress may also damage mitochondria, initiating the release of ROS or mtDNA, which further promotes a pro-inflammatory response via inflammasome activation and cytokine production (Riley and Tait, 2020; Zhou et al., 2011). We examined cytokine levels in both the PFC and plasma to determine whether our gestational stress protocol induced inflammation in association with mitochondrial dysfunction. In the PFC, stress significantly increased levels of TNF-α and these levels negatively correlated with PFC mitochondrial respiration. In the periphery, parity increased levels of TNF-α, IL6, IL-1β, IL-18, IFN-γ, and VEGF. Additionally, plasma TNF-α, but not other measured cytokines similarly correlated with PFC mitochondrial respiration. Despite the increased levels of TNF-α in both plasma and PFC and their significant correlations with PFC mitochondrial respiration, there was no significant correlation between neural and systemic TNF-α. This finding replicates previous studies reporting a lack of correlation between central and peripheral immune activation (Boufidou et al., 2009; Miller et al., 2019) and further highlights the unique and complex nature of postpartum inflammation. Indeed, this complexity is further underlined by the conflicting reports on gestational stress-induced inflammation at both the central and peripheral levels in the rodent literature (Dye et al., 2022), possibly due to differences in the timing of measurement and stress protocols. Measurements taken at GD21 identified increased levels of IL-1β and Interferon gamma (IFN-γ) in the PFC of gestationally stressed females, but no changes in plasma cytokine levels from non-stressed pregnant counterparts (Lenz et al., 2019). A separate study found increased IL-6 and decreased IL-1β in the PFC in gestationally stressed postpartum rats the day after parturition but did not measure in the periphery. One month postpartum, O’Mahoney and colleagues observed increased TNF-α, IL-1β, and IL-10 levels in the plasma of gestationally stressed rats but did not observe any differences in the PFC (O’Mahony et al., 2006). These studies together with our findings here suggest a potential timeline of stress-induced neuroimmune changes during gestation that possibly develop into systemic inflammation in the postpartum. Longitudinal sampling will be necessary to fully establish the sequence of events surrounding stress-evoked increases in inflammation. Studies in patients, while limited, are slightly more consistent, with PPD women exhibiting enhanced TNF-α both systemically (Boufidou et al., 2009; Bränn et al., 2018) and centrally (in cerebrospinal fluid) and these levels correlating with depressive symptoms (Boufidou et al., 2009). Moreover, levels of serum IL-6 were increased in postpartum women with increased depression scores relative to prenatal time points (Maes et al., 2000) and serum IL-18 was increased in women 8 weeks postpartum (Bränn et al., 2020). Our cytokine data, coupled with these studies, lend further support to TNF-α as a cytokine that warrants further examination.

Importantly, our findings reveal a relationship between PFC mitochondrial function, inflammation, and PPD-relevant behaviors. We identified significant negative correlations between PFC mitochondrial respiration and the latency to immobility in the forced swim test, as well as between PFC mitochondrial respiration and TNF-α levels in the PFC and plasma. While the nature of our study does not allow us to draw conclusions on causality or directionality, it opens the door to several possibilities. One is that gestational stress has parallel effects on mitochondria, behavior, and inflammation. Another is that gestational stress induces inflammation that induces mitochondrial and behavioral dysfunction. The more tantalizing possibility, however, is that gestational stress induces PPD-relevant behaviors and inflammation via effects on brain mitochondrial function. We have previously shown that mitochondrial function in specific brain regions can drive behavioral responses (Hollis et al., 2018, 2015; van der Kooij et al., 2018), so it is plausible that mitochondria act as mediators to link stress with behavioral consequences. Moreover, mitochondria have been recently shown to activate inflammasomes and trigger the release of pro-inflammatory cytokines (West, 2017). The significant relationship between plasma cytokine levels and PFC mitochondrial respiration is particularly exciting as it suggests that plasma TNF-α may act as a biomarker for central mitochondrial function, and possibly, PPD. Further studies will be necessary to fully characterize the biomarker potential of TNF-α for PFC mitochondrial function and PPD. As cytokines can also act to disrupt mitochondrial function (Hollis et al., 2022) future studies that manipulate mitochondrial function will be needed to determine the precise role of PFC mitochondrial respiration in PPD-relevant behaviors and inflammation.

It is important to note that our mitochondrial measures were performed at a single mid-postpartum time point. Thus, we cannot rule out potential effects of dam-offspring interactions that may elicit neuronal changes following gestational stress (Baker et al., 2008; Champagne and Meaney, 2006a; Leuner et al., 2014b; Smith et al., 2004), nor potential interactions with hormones stimulating lactation, particularly as we observed differences in maternal nursing and grooming behaviors. Moreover, our study does not identify the precise time point at which mitochondrial respiration was affected. We do not know whether mitochondrial alterations occurred following parturition or developed across gestation with successive stress exposures. Nor do we know how long these changes persist in the postpartum period. Follow-up experiments that identify the timing, duration, and specificity of mitochondrial alterations will be necessary to clarify these points. Despite these limitations, our study represents an important first step in establishing the effects of gestational stress and pregnancy to alter postpartum brain mitochondrial function.

Since mitochondria are involved in many biological processes, the mechanisms contributing to our observed results could be multi-faceted. Importantly, in the current study, mitochondrial dysfunction was only observed when both stress and pregnancy were involved, pointing to interactions between the HPA and HPG axes. Pregnancy and parturition are marked by dramatic fluctuations in reproductive hormones, such as estrogen that decreases drastically in the postpartum (Brummelte and Galea, 2016). 17*β**-***Estradiol (E_2_), has been shown to benefit mitochondrial function by reducing oxidative damage and promoting energy production via oxidative phosphorylation (OXPHOS) (Brinton, 2008b), and E_2_ affects stress responses of the hypothalamic-pituitary-adrenal (HPA) axis (Handa et al., 2012). Estrogen receptors can localize to mitochondria, where their actions can affect respiratory activity, mitochondrial enzymatic activity, and reduce lipid peroxidation (Irwin et al., 2012; Yang et al., 2009, 2004). It is possible that the fluctuation of estrogen levels during and shortly following pregnancy have downstream effects on mitochondrial function, promoting a dysfunctional phenotype. Similarly, corticosterone (CORT) may also have a mechanistic role, considering that fluctuations in this hormone also occur during pregnancy (Brummelte and Galea, 2016) and glucocorticoid receptors can localize to mitochondria and regulate their function (Du et al., 2009). While these hypotheses will provide a crucial stepping-off point to future research, there are certainly other mechanisms to consider and the possibility of a combination of these processes should not be ruled out.

## 5. Conclusion

Despite the multitude of connections between mitochondrial function and proposed mechanisms of PPD, to our knowledge, this is the first study to directly examine brain mitochondrial function in pregnancy or PPD. Overall, our results suggest that mitochondria could act as an upstream regulator of stress-induced processes which affect behavioral output. However, future studies to identify potential downstream targets, or whether dysregulation is occurring upstream that has direct consequences on mitochondrial function, are required to clarify the mechanisms involved. Finally, experiments in which mitochondrial function is either enhanced or reduced (*e.g.,* with treatments of an antioxidant or an electron transport system complex inhibitor, respectively) would provide necessary information on the direct role of these organelles on the manifestation of postpartum depressive-like behaviors. With these aims in mind, mitochondria could be a potential therapeutic target in the development of more effective and cost-efficient treatment for PPD patients, specifically for pregnant individuals who are more vulnerable to the combined effects of stress and pregnancy.

## Funding Support

This work was supported by the Department of Veteran Affairs through VISN7 Research Development Award; as well as the NIH with a COBRE Target Faculty award (P20GM109091) to FH and RO1DK132948 (R.C.W.); and USC institutional funds via a UofSC ASPIRE I award and institutional start-up funding to FH.

## CRediT authorship contribution statement

**Erin Gorman-Sandler:** Conceptualization, Investigation, Formal analysis, Data Curation, Writing – original draft, Writing – review and editing. **Breanna Robertson:** Investigation, Formal analysis, Data Curation. **Jesseca Crawford:** Investigation. **Olufunke A Arishe:** Investigation. **R. Clinton Webb:** Writing – review and editing. **Fiona Hollis:** Conceptualization, Formal Analysis, Data Curation, Writing – original draft, Writing – review and editing, Funding Acquisition.

## Declaration of competing interest

The authors have no interests to declare.

## Acknowledgements

The authors would like to thank Ms. Hannah Burzynski and Dr. Lawrence Reagan for training and assistance with the BioPlex assays. Additionally, we would like to thank Dr. Lawrence Reagan for helpful discussions and manuscript feedback.

## Supplementary Figure Legends

**Supplementary Figure 1. Main effects of parity in water baseline and no effects of stress on 0.25% sucrose preference or total liquid consumption. (A)** Water intake baseline during the week prior to parturition exhibited a significant main effect of parity (*two-way ANOVA;* parity: F_(1,29)_ = 15.53; p = 0.0005) but no significant interaction or main effect of stress (interaction: F_(1,29)_ = 1.516; p = 0.2282, stress: F_(1,29)_ = 0.0046; p = 0.9462) such that primiparous females had significantly higher water intake compared to nulliparous groups (*Dunnett’s multiple comparisons test*; C vs. S: p = 0.6711, C vs. P: p = 0.0031, C vs. P+S: p = 0.0302). n = 6-11/group. **(B)** Preference for a weak (0.25%) sucrose solution exhibited trends for a main interaction between stress and parity and parity, but no significant main effect of stress (*two-way ANOVA;* interaction: F_(1,19)_ = 3.324; p = 0.084, parity: F_(1,19)_ = 3.862; p = 0.0642, stress: F_(1,19) = 0.1648; p = 0.6893_). n = 5-6/group. **(C)** Total fluid intake during 1% sucrose measurements were not significantly different between groups, evidenced by no main effects of parity, stress, nor interaction (*two-way ANOVA;* interaction: F_(1,19)_ = 0.3717; p = 0.5493, parity: F_(1,19)_ = 1.288; p = 0.2705, stress: F_(1,19) = 0.0524; p = 0.8214_). n = 5-6/group. All data are represented as mean ± SEM. *p≤0.05, **p≤0.01,***p≤0.001

**Supplementary Figure 2. No significant differences in open arm entries, distance, or velocity during the Elevated Plus Maze. (A)** Open arm entries were not different between groups as evidenced by no main effects of parity, stress, nor interaction (*two-way ANOVA;* interaction: F_(1,35)_ = 2.712; p = 0.1085, parity: F_(1,35)_ = 0.5204; p = 0.4755, stress: F_(1,35) = 1.757; p= 0.1936_). n = 9-11/group. **(B)** All groups traveled similar distances during testing, with no significant main effects of parity, stress, nor interaction (*two-way ANOVA;* interaction: F_(1,35)_ = 0.2479; p = 0.6217, parity: F_(1,35)_ = 0.6646; p = 0.4204, stress: F_(1,35) = 0.7977; p = 0.3779_). n = 9-11/group. **(C)** Average velocity was similar between groups (*two-way ANOVA;* interaction: F_(1,35)_ = 0.2603; p = 0.6131, parity: F_(1,35)_ = 0.5009; p = 0.4838, stress: F_(1,35) = 0.8196; p = 0.3715_). n = 9-11/group. All data are represented as mean ± SEM.

**Supplementary Figure 3. No effects of stress or parity on mitochondrial content in PFC or NAc. (A)** There were no significant differences between groups in expression of the mitochondrial outer membrane protein, TOM20 in the PFC *(two-way ANOVA;* interaction: F_(1,33)_ = 1.73; p = 0.1975, parity: F_(1,33)_ = 0.0894; p = 0.7669, stress: F_(1,33)_ = 0.0497; p = 0.8249). n = 7-11/group. **(B)** There were no significant differences in complex I and II-linked ETS measures in the NAc between groups (*two-way ANOVA;* interaction: F_(1,36)_ = 0.1463; p = 0.7044, parity: F_(1,36)_ = 0.054; p = 0.8176, stress: F_(1,36)_ = 0.2185; p = 0.643). n=9-11/group. All data are represented as mean ± SEM.

**Supplementary Figure 4. There were no effects of parity or gestational stress on other measured PFC cytokine levels.** There were no significant main interactions, or effects of stress or parity on levels of **(A)** IL-1β (*two-way ANOVA*; interaction: F_(1,32)_ = 0.0537; p = 0.8182, parity: F_(1,32)_ = 2.961; p = 0.095, stress: F_(1,32)_ = 0.0524; p = 0.8203). n = 8-10/group; **(B)** IL-1a (*two-way ANOVA*; interaction: F_(1,35)_ = 0.6768; p = 0.4162, parity: F_(1,35)_ = 0.1233; p = 0.7275, stress: F_(1,35)_ = 0.2658; p = 0.6094). n = 9-11/group; **(C)** IL-6 (*two-way ANOVA*; interaction: F_(1,25)_ = 0.1604; p = 0.6922, parity: F_(1,25)_ = 0.6928; p = 0.4131, stress: F_(1,25)_ = 1.719; p = 0.2018). n = 7-8/group; **(D)** IL-10 (*two-way ANOVA*; interaction: F_(1,29)_ = 0.7102; p = 0.4063, parity: F_(1,29)_ = 0.1832; p = 0.6718, stress: F_(1,29)_ = 1.788; p = 0.1916). n = 7-10/group. All data are represented as mean ± SEM.

**Supplementary Figure 5. Significant effect of parity on additional plasma cytokine levels. (A)** IL-1β levels exhibited a significant main effect of parity (*two-way ANOVA*; interaction: F_(1,35)_ = 0.8624; p = 0.3594, parity: F_(1,35)_ = 11.26; p = 0.0019, stress: F_(1,35)_ = 0.7097; p = 0.4053), but *posthoc* analysis revealed no significant differences between groups. (*Dunnett’s multiple comparisons test*; C vs. S: p = 0.4151, C vs. P: p = 0.2396, C vs. P+S: p = 0.1937). n = 8-11/group. **(B)** Analysis of IL-6 revealed a significant main effect of parity (*two-way ANOVA*; interaction: F_(1,34)_ = 1.527; p = 0.225, parity: F_(1,34)_ = 9.997; p = 0.0033, stress: F_(1,34)_ = 0.052; p = 0.8209) and trends for P+S females to have higher levels compared to control (*Dunnett’s multiple comparisons test*; C vs. S: p = 0.8018, C vs. P: p = 0.4098, C vs. P+S: p = 0.0505).n = 8-11/group. **(C)** There was a trend for an interaction between parity and stress and a significant main effect of parity but no main effect of stress in IL-18 levels (*two-way ANOVA*; interaction: F_(1,35)_ = 3.632; p = 0.0649, parity: F_(1,35)_ = 7.943; p = 0.0079, stress: F_(1,35)_ = 0.5997; p = 0.4439). *Posthoc* analysis revealed no significant differences between groups (*Dunnett’s multiple comparisons test*; C vs. S: p = 0.1259, C vs. P: p = 0.8673, C vs. P+S: p = 0.343). n = 8-11/group. **(D)** Similarly, there was a trend for an interaction and a significant main effect of parity in IFN-γ levels (*two-way ANOVA*; interaction: F_(1,34)_ = 2.933; p = 0.0959, parity: F_(1,34)_ = 11.33; p = 0.0019, stress: F_(1,34)_ = 0.7511; p = 0.3922). *Posthoc* analysis revealed a significant difference between P+S and control such that P+S females had higher levels (*Dunnett’s multiple comparisons test*; C vs. S: p = 0.897, C vs. P: p = 0.5829, C vs. P+S: p = 0.0123). n = 8-11/group. **(E)** There was a significant main effect of parity, but no effect of stress or interaction in VEGF (*two-way ANOVA*; interaction: F_(1,35)_ = 0.5525; p = 0.4622, parity: F_(1,35)_ = 7.403; p = 0.0101, stress: F_(1,35)_ = 0.388; p = 0.5374). *Posthoc* analysis revealed no significant differences between groups (*Dunnett’s multiple comparisons test*; C vs. S: p = 0.6145, C vs. P: p = 0.3955, C vs. P+S: p = 0.3228). n = 8-11/group. **(F)** TNFα in PFC does not correlate with TNFα in plasma (*Simple linear regression*; p = 0.6132, R^2^ = 0.0076). n = 37. Nulliparous controls =C, nulliparous stressed =S, primiparous controls =P, and primiparous stressed =P+S. All data are represented as mean ± SEM. *p≤0.05, **p≤0.01, ***p≤0.001, ****p≤0.0001.

Conflict of Interest

## Declaration of competing interest

The authors have no competing interests or conflict to declare.

## Notes

### Competing Interest Statement

The authors have declared no competing interest.

## References

Accardi, M.V., Daniels, B.A., Brown, P.M.G.E., Fritschy, J.-M., Tyagarajan, S.K., Bowie, D., 2014. Mitochondrial reactive oxygen species regulate the strength of inhibitory GABA-mediated synaptic transmission. Nat. Commun. 5, 3168. https://doi.org/10.1038/ncomms4168

Allen, J., Caruncho, H.J., Kalynchuk, L.E., 2021. Severe life stress, mitochondrial dysfunction, and depressive behavior: A pathophysiological and therapeutic perspective. Mitochondrion 56, 111–117. https://doi.org/10.1016/j.mito.2020.11.010

Anderson, G., Maes, M., 2013. Postpartum depression: psychoneuroimmunological underpinnings and treatment. Neuropsychiatr. Dis. Treat. 9, 277–287. https://doi.org/10.2147/NDT.S25320

Austin, M.-P.V., Hadzi-Pavlovic, D., Priest, S.R., Reilly, N., Wilhelm, K., Saint, K., Parker, G., 2010. Depressive and anxiety disorders in the postpartum period: how prevalent are they and can we improve their detection? Arch. Womens Ment. Health 13, 395–401. https://doi.org/10.1007/s00737-010-0153-7

Baker, S., Chebli, M., Rees, S., LeMarec, N., Godbout, R., Bielajew, C., 2008. Effects of gestational stress: 1. Evaluation of maternal and juvenile offspring behavior. Brain Res. 1213, 98–110. https://doi.org/10.1016/j.brainres.2008.03.035

Baker, S.L., Kentner, A.C., Konkle, A.T.M., Santa-Maria Barbagallo, L., Bielajew, C., 2006. Behavioral and physiological effects of chronic mild stress in female rats. Physiol. Behav. 87, 314–322. https://doi.org/10.1016/j.physbeh.2005.10.019

Bali, A., Jaggi, A.S., 2014. Multifunctional aspects of allopregnanolone in stress and related disorders. Prog. Neuropsychopharmacol. Biol. Psychiatry 48, 64–78. https://doi.org/10.1016/j.pnpbp.2013.09.005

Becker, M., Weinberger, T., Chandy, A., Schmukler, S., 2016. Depression During Pregnancy and Postpartum. Curr. Psychiatry Rep. 18, 32. https://doi.org/10.1007/s11920-016-0664-7

Bolaños, C.A., Willey, M.D., Maffeo, M.L., Powers, K.D., Kinka, D.W., Grausam, K.B., Henderson, R.P., 2008. Antidepressant Treatment Can Normalize Adult Behavioral Deficits Induced by Early-Life Exposure to Methylphenidate. Biol. Psychiatry, Impulse Control: Aggression, Addiction, and Attention Deficits 63, 309–316. https://doi.org/10.1016/j.biopsych.2007.06.024

Bordt, E.A., Polster, B.M., 2014. NADPH oxidase- and mitochondria-derived reactive oxygen species in proinflammatory microglial activation: a bipartisan affair? Free Radic. Biol. Med. 76, 34–46. https://doi.org/10.1016/j.freeradbiomed.2014.07.033

Bordt, E.A., Smith, C.J., Demarest, T.G., Bilbo, S.D., Kingsbury, M.A., 2019. Mitochondria, Oxytocin, and Vasopressin: Unfolding the inflammatory protein response. Neurotox. Res. 36, 239–256. https://doi.org/10.1007/s12640-018-9962-7

Boufidou, F., Lambrinoudaki, I., Argeitis, J., Zervas, I.M., Pliatsika, P., Leonardou, A.A., Petropoulos, G., Hasiakos, D., Papadias, K., Nikolaou, C., 2009. CSF and plasma cytokines at delivery and postpartum mood disturbances. J. Affect. Disord. 115, 287– 292. https://doi.org/10.1016/j.jad.2008.07.008

Bränn, E., Edvinsson, Å., Rostedt Punga, A., Sundström-Poromaa, I., Skalkidou, A., 2019. Inflammatory and anti-inflammatory markers in plasma: from late pregnancy to early postpartum. Sci. Rep. 9, 1863. https://doi.org/10.1038/s41598-018-38304-w

Bränn, E., Fransson, E., White, R.A., Papadopoulos, F.C., Edvinsson, Å., Kamali-Moghaddam, M., Cunningham, J.L., Sundström-Poromaa, I., Skalkidou, A., 2020. Inflammatory markers in women with postpartum depressive symptoms. J. Neurosci. Res. 98, 1309– 1321.

Brinton, R.D., 2008a. The healthy cell bias of estrogen action: mitochondrial bioenergetics and neurological implications. Trends Neurosci. 31, 529–537. https://doi.org/10.1016/j.tins.2008.07.003

Brinton, R.D., 2008b. The healthy cell bias of estrogen action: mitochondrial bioenergetics and neurological implications. Trends Neurosci. 31, 529–537. https://doi.org/10.1016/j.tins.2008.07.003

Brummelte, S., Galea, L.A.M., 2016. Postpartum depression: Etiology, treatment and consequences for maternal care. Horm. Behav., Parental Care 77, 153–166. https://doi.org/10.1016/j.yhbeh.2015.08.008

Brummelte, S., Galea, L.A.M., 2010. Chronic corticosterone during pregnancy and postpartum affects maternal care, cell proliferation and depressive-like behavior in the dam. Horm. Behav. 58, 769–779. https://doi.org/10.1016/j.yhbeh.2010.07.012

Brunton, P.J., Russell, J.A., Hirst, J.J., 2014. Allopregnanolone in the brain: protecting pregnancy and birth outcomes. Prog. Neurobiol. 113, 106–136. https://doi.org/10.1016/j.pneurobio.2013.08.005

Burtscher, J., Zangrandi, L., Schwarzer, C., Gnaiger, E., 2015. Differences in mitochondrial function in homogenated samples from healthy and epileptic specific brain tissues revealed by high-resolution respirometry. Mitochondrion 25, 104–112. https://doi.org/10.1016/j.mito.2015.10.007

Camacho-Arroyo, I., González-Arenas, A., Jiménez-Arellano, C., Morimoto, S., Galván-Rosas, A., Gómora-Arrati, P., García-Juárez, M., González-Flores, O., 2018. Sex hormone levels and expression of their receptors in lactating and lactating pregnant rats. J. Steroid Biochem. Mol. Biol. 178, 213–220. https://doi.org/10.1016/j.jsbmb.2017.12.015

Champagne, F.A., 2004. Variations in Nucleus Accumbens Dopamine Associated with Individual Differences in Maternal Behavior in the Rat. J. Neurosci. 24, 4113–4123. https://doi.org/10.1523/JNEUROSCI.5322-03.2004

Champagne, F.A., Meaney, M.J., 2006a. Stress During Gestation Alters Postpartum Maternal Care and the Development of the Offspring in a Rodent Model. Biol. Psychiatry 59, 1227–1235. https://doi.org/10.1016/j.biopsych.2005.10.016

Champagne, F.A., Meaney, M.J., 2006b. Stress during gestation alters postpartum maternal care and the development of the offspring in a rodent model. Biol. Psychiatry 59, 1227– 1235. https://doi.org/10.1016/j.biopsych.2005.10.016

Cheng, A., Hou, Y., Mattson, M.P., 2010. Mitochondria and neuroplasticity. ASN NEURO 2, e00045. https://doi.org/10.1042/AN20100019

Christian, L.M., 2015. Stress and Immune Function during Pregnancy: An Emerging Focus in Mind-Body Medicine. Curr. Dir. Psychol. Sci. 24, 3–9. https://doi.org/10.1177/0963721414550704

Corwin, E.J., Pajer, K., Paul, S., Lowe, N., Weber, M., McCarthy, D.O., 2015. Bidirectional psychoneuroimmune interactions in the early postpartum period influence risk of postpartum depression. Brain. Behav. Immun. 49, 86–93. https://doi.org/10.1016/j.bbi.2015.04.012

Csányi, A., Bóta, J., Falkay, G., Gáspár, R., Ducza, E., 2016. The Effects of Female Sexual Hormones on the Expression of Aquaporin 5 in the Late-Pregnant Rat Uterus. Int. J. Mol. Sci. 17, 1300. https://doi.org/10.3390/ijms17081300

Darnaudéry, M., Dutriez, I., Viltart, O., Morley-Fletcher, S., Maccari, S., 2004. Stress during gestation induces lasting effects on emotional reactivity of the dam rat. Behav. Brain Res. 153, 211–216. https://doi.org/10.1016/j.bbr.2003.12.001

Depression Among Women | Depression | Reproductive Health | CDC [WWW Document], 2020. URL https://www.cdc.gov/reproductivehealth/depression/index.htm (accessed 4.20.21).

Drevets, W.C., Price, J.L., Furey, M.L., 2008. Brain structural and functional abnormalities in mood disorders: implications for neurocircuitry models of depression. Brain Struct. Funct. 213, 93–118. https://doi.org/10.1007/s00429-008-0189-x

Du, J., Wang, Y., Hunter, R., Wei, Y., Blumenthal, R., Falke, C., Khairova, R., Zhou, R., Yuan, P., Machado-Vieira, R., McEwen, B.S., Manji, H.K., 2009. Dynamic regulation of mitochondrial function by glucocorticoids. Proc. Natl. Acad. Sci. U. S. A. 106, 3543– 3548. https://doi.org/10.1073/pnas.0812671106

Du Preez, A., Eum, J., Eiben, I., Eiben, P., Zunszain, P.A., Pariante, C.M., Thuret, S., Fernandes, C., 2021. Do different types of stress differentially alter behavioural and neurobiological outcomes associated with depression in rodent models? A systematic review. Front. Neuroendocrinol. 61, 100896. https://doi.org/10.1016/j.yfrne.2020.100896

Duman, R.S., 2014. Neurobiology of stress, depression, and rapid acting antidepressants: remodeling synaptic connections. Depress. Anxiety 31, 291–296.

Dye, C., Lenz, K.M., Leuner, B., 2022. Immune System Alterations and Postpartum Mental Illness: Evidence From Basic and Clinical Research. Front. Glob. Womens Health 2.

Feldthusen, A.-D., Larsen, J., Pedersen, P.L., Toft Kristensen, T., Kvetny, J., 2014. Pregnancy-induced alterations in mitochondrial function in euthyroid pregnant women and pregnant women with subclinical hypothyroidism; relation to adverse outcome. J. Clin. Transl. Endocrinol. 1, e13–e17. https://doi.org/10.1016/j.jcte.2013.12.003

Filiou, M.D., Sandi, C., 2019. Anxiety and Brain Mitochondria: A Bidirectional Crosstalk. Trends Neurosci. 42, 573–588. https://doi.org/10.1016/j.tins.2019.07.002

Filiou, M.D., Zhang, Y., Teplytska, L., Reckow, S., Gormanns, P., Maccarrone, G., Frank, E., Kessler, M.S., Hambsch, B., Nussbaumer, M., Bunck, M., Ludwig, T., Yassouridis, A., Holsboer, F., Landgraf, R., Turck, C.W., 2011. Proteomics and metabolomics analysis of a trait anxiety mouse model reveals divergent mitochondrial pathways. Biol. Psychiatry 70, 1074–1082. https://doi.org/10.1016/j.biopsych.2011.06.009

Franceschelli, A., Herchick, S., Thelen, C., Papadopoulou-Daifoti, Z., Pitychoutis, P.M., 2014. Sex differences in the chronic mild stress model of depression. Behav. Pharmacol. 25, 372–383. https://doi.org/10.1097/FBP.0000000000000062

Frisbee, J.C., Brooks, S.D., Stanley, S.C., d’Audiffret, A.C., 2015. An Unpredictable Chronic Mild Stress Protocol for Instigating Depressive Symptoms, Behavioral Changes and Negative Health Outcomes in Rodents. J. Vis. Exp. JoVE 53109. https://doi.org/10.3791/53109

Gemmel, M., Harmeyer, D., Bögi, E., Fillet, M., Hill, L.A., Hammond, G.L., Charlier, T.D., Pawluski, J.L., 2018. Perinatal fluoxetine increases hippocampal neurogenesis and reverses the lasting effects of pre-gestational stress on serum corticosterone, but not on maternal behavior, in the rat dam. Behav. Brain Res. 339, 222–231. https://doi.org/10.1016/j.bbr.2017.11.038

Ghosal, S., Hare, B., Duman, R.S., 2017. Prefrontal Cortex GABAergic Deficits and Circuit Dysfunction in the Pathophysiology and Treatment of Chronic Stress and Depression. Curr. Opin. Behav. Sci. 14, 1–8. https://doi.org/10.1016/j.cobeha.2016.09.012

Haim, A., Albin-Brooks, C., Sherer, M., Mills, E., Leuner, B., 2016. The effects of gestational stress and SSRI antidepressant treatment on structural plasticity in the postpartum brain - a translational model for postpartum depression. Horm. Behav. 77, 124–131. https://doi.org/10.1016/j.yhbeh.2015.05.005

Haim, A., Sherer, M., Leuner, B., 2014. Gestational stress induces persistent depressive-like behavior and structural modifications within the postpartum nucleus accumbens. Eur. J. Neurosci. 40, 3766–3773. https://doi.org/10.1111/ejn.12752

Handa, R.J., Mani, S.K., Uht, R.M., 2012. Estrogen receptors and the regulation of neural stress responses. Neuroendocrinology 96, 111–118. https://doi.org/10.1159/000338397

Hillerer, K.M., Jacobs, V.R., Fischer, T., Aigner, L., 2014. The Maternal Brain: An Organ with Peripartal Plasticity. Neural Plast. 2014, 574159. https://doi.org/10.1155/2014/574159

Hillerer, K.M., Reber, S.O., Neumann, I.D., Slattery, D.A., 2011. Exposure to chronic pregnancy stress reverses peripartum-associated adaptations: implications for postpartum anxiety and mood disorders. Endocrinology 152, 3930–3940. https://doi.org/10.1210/en.2011-1091

Hoekzema, E., Barba-Müller, E., Pozzobon, C., Picado, M., Lucco, F., García-García, D., Soliva, J.C., Tobeña, A., Desco, M., Crone, E.A., Ballesteros, A., Carmona, S., Vilarroya, O., 2017. Pregnancy leads to long-lasting changes in human brain structure. Nat. Neurosci. 20, 287–296. https://doi.org/10.1038/nn.4458

Hollis, F., Duclot, F., Gunjan, A., Kabbaj, M., 2011. Individual differences in the effect of social defeat on anhedonia and histone acetylation in the rat hippocampus. Horm. Behav. 59, 331–337. https://doi.org/10.1016/j.yhbeh.2010.09.005

Hollis, F., Kooij, M.A. van der, Zanoletti, O., Lozano, L., Cantó, C., Sandi, C., 2015. Mitochondrial function in the brain links anxiety with social subordination. Proc. Natl. Acad. Sci. 112, 15486–15491. https://doi.org/10.1073/pnas.1512653112

Hollis, F., Mitchell, E.S., Canto, C., Wang, D., Sandi, C., 2018. Medium chain triglyceride diet reduces anxiety-like behaviors and enhances social competitiveness in rats. Neuropharmacology 138, 245–256. https://doi.org/10.1016/j.neuropharm.2018.06.017

Hollis, F., Pope, B.S., Gorman-Sandler, E., Wood, S.K., 2022. Neuroinflammation and Mitochondrial Dysfunction Link Social Stress to Depression. Curr. Top. Behav. Neurosci. 54, 59–93. https://doi.org/10.1007/7854_2021_300

Hollis, F., Wang, H., Dietz, D., Gunjan, A., Kabbaj, M., 2010. The effects of repeated social defeat on long-term depressive-like behavior and short-term histone modifications in the hippocampus in male Sprague–Dawley rats. Psychopharmacology (Berl.) 211, 69–77. https://doi.org/10.1007/s00213-010-1869-9

Irwin, R.W., Yao, J., To, J., Hamilton, R.T., Cadenas, E., Brinton, R.D., 2012. Selective oestrogen receptor modulators differentially potentiate brain mitochondrial function. J. Neuroendocrinol. 24, 236–248. https://doi.org/10.1111/j.1365-2826.2011.02251.x

Jolley, S.N., Elmore, S., Barnard, K.E., Carr, D.B., 2007. Dysregulation of the hypothalamic-pituitary-adrenal axis in postpartum depression. Biol. Res. Nurs. 8, 210–222. https://doi.org/10.1177/1099800406294598

Kanes, S., Colquhoun, H., Gunduz-Bruce, H., Raines, S., Arnold, R., Schacterle, A., Doherty, J., Epperson, C.N., Deligiannidis, K.M., Riesenberg, R., Hoffmann, E., Rubinow, D., Jonas, J., Paul, S., Meltzer-Brody, S., 2017. Brexanolone (SAGE-547 injection) in post-partum depression: a randomised controlled trial. Lancet Lond. Engl. 390, 480–489. https://doi.org/10.1016/S0140-6736(17)31264-3

Lancaster, C.A., Gold, K.J., Flynn, H.A., Yoo, H., Marcus, S.M., Davis, M.M., 2010. Risk factors for depressive symptoms during pregnancy: a systematic review. Am. J. Obstet. Gynecol. 202, 5–14. https://doi.org/10.1016/j.ajog.2009.09.007

Lenz, K.M., Post, C., Castaneda, A.J., Banta, P., Nelson, L.H., Saulsbery, A.I., Leuner, B., 2019. Abstract # 3185 Central immune alterations in a gestational stress animal model of postpartum depression. Brain. Behav. Immun., PsychoNeuroImmunology Research Society’s 25th Annual Scientific Meeting 76, e38. https://doi.org/10.1016/j.bbi.2018.11.295

Leuner, B., Fredericks, P.J., Nealer, C., Albin-Brooks, C., 2014a. Chronic Gestational Stress Leads to Depressive-Like Behavior and Compromises Medial Prefrontal Cortex Structure and Function during the Postpartum Period. PLOS ONE 9, e89912. https://doi.org/10.1371/journal.pone.0089912

Leuner, B., Fredericks, P.J., Nealer, C., Albin-Brooks, C., 2014b. Chronic Gestational Stress Leads to Depressive-Like Behavior and Compromises Medial Prefrontal Cortex Structure and Function during the Postpartum Period. PLoS ONE 9, e89912. https://doi.org/10.1371/journal.pone.0089912

Leuner, B., Gould, E., 2010. Dendritic Growth in Medial Prefrontal Cortex and Cognitive Flexibility Are Enhanced during the Postpartum Period. J. Neurosci. 30, 13499–13503. https://doi.org/10.1523/JNEUROSCI.3388-10.2010

Li, Z., Okamoto, K.-I., Hayashi, Y., Sheng, M., 2004. The importance of dendritic mitochondria in the morphogenesis and plasticity of spines and synapses. Cell 119, 873–887.

Liang, J.J., Rasmusson, A.M., 2018. Overview of the Molecular Steps in Steroidogenesis of the GABAergic Neurosteroids Allopregnanolone and Pregnanolone. Chronic Stress 2, 2470547018818555. https://doi.org/10.1177/2470547018818555

Lonstein, J.S., 2007. Regulation of anxiety during the postpartum period. Front. Neuroendocrinol. 28, 115–141. https://doi.org/10.1016/j.yfrne.2007.05.002

Madrigal, J.L., Olivenza, R., Moro, M.A., Lizasoain, I., Lorenzo, P., Rodrigo, J., Leza, J.C., 2001. Glutathione depletion, lipid peroxidation and mitochondrial dysfunction are induced by chronic stress in rat brain. Neuropsychopharmacol. Off. Publ. Am. Coll. Neuropsychopharmacol. 24, 420–429. https://doi.org/10.1016/S0893-133X(00)00208-6

Maguire, J., Mody, I., 2008. GABAAR Plasticity during Pregnancy: Relevance to Postpartum Depression. Neuron 59, 207–213. https://doi.org/10.1016/j.neuron.2008.06.019

McEwen, A.M., Burgess, D.T.A., Hanstock, C.C., Seres, P., Khalili, P., Newman, S.C., Baker, G.B., Mitchell, N.D., Khudabux-Der, J., Allen, P.S., LeMelledo, J.-M., 2012. Increased Glutamate Levels in the Medial Prefrontal Cortex in Patients with Postpartum Depression. Neuropsychopharmacology 37, 2428–2435. https://doi.org/10.1038/npp.2012.101

Melón, L., Hammond, R., Lewis, M., Maguire, J., 2018. A Novel, Synthetic, Neuroactive Steroid Is Effective at Decreasing Depression-Like Behaviors and Improving Maternal Care in Preclinical Models of Postpartum Depression. Front. Endocrinol. 9, 703. https://doi.org/10.3389/fendo.2018.00703

Meltzer-Brody, S., Kanes, S.J., 2020. Allopregnanolone in postpartum depression: Role in pathophysiology and treatment. Neurobiol. Stress 12, 100212. https://doi.org/10.1016/j.ynstr.2020.100212

Miller, E.S., Sakowicz, A., Roy, A., Yang, A., Sullivan, J.T., Grobman, W.A., Wisner, K.L., 2019. Plasma and cerebrospinal fluid inflammatory cytokines in perinatal depression. Am. J. Obstet. Gynecol. 220, 271.e1–271.e10. https://doi.org/10.1016/j.ajog.2018.12.015

Molendijk, M.L., de Kloet, E.R., 2015. Immobility in the forced swim test is adaptive and does not reflect depression. Psychoneuroendocrinology 62, 389–391. https://doi.org/10.1016/j.psyneuen.2015.08.028

Morava, É., Kozicz, T., 2013. Mitochondria and the economy of stress (mal)adaptation. Neurosci. Biobehav. Rev. 37, 668–680. https://doi.org/10.1016/j.neubiorev.2013.02.005

Moses-Kolko, E.L., Perlman, S.B., Wisner, K.L., James, J., Saul, A.T., Phillips, M.L., 2010. Abnormally Reduced Dorsomedial Prefrontal Cortical Activity and Effective Connectivity With Amygdala in Response to Negative Emotional Faces in Postpartum Depression. Am. J. Psychiatry 167, 1373–1380. https://doi.org/10.1176/appi.ajp.2010.09081235

Musatov, A., Robinson, N.C., 2012. Susceptibility of mitochondrial electron-transport complexes to oxidative damage. Focus on cytochrome c oxidase. Free Radic. Res. 46, 1313–1326. https://doi.org/10.3109/10715762.2012.717273

O’Hara, M.W., 2009. Postpartum depression: what we know. J. Clin. Psychol. 65, 1258–1269. https://doi.org/10.1002/jclp.20644

O’hara, M.W., Swain, A.M., 1996. Rates and risk of postpartum depression—a meta-analysis. Int. Rev. Psychiatry 8, 37–54. https://doi.org/10.3109/09540269609037816

O’Hara, M.W., Wisner, K.L., 2014. Perinatal mental illness: Definition, description and aetiology. Best Pract. Res. Clin. Obstet. Gynaecol., Perinatal Mental Health: Guidance for the Obstetrician-Gynaecologist 28, 3–12. https://doi.org/10.1016/j.bpobgyn.2013.09.002

O’Mahony, S.M., Myint, A.-M., van den Hove, D., Desbonnet, L., Steinbusch, H., Leonard, B.E., 2006. Gestational stress leads to depressive-like behavioural and immunological changes in the rat. Neuroimmunomodulation 13, 82–88. https://doi.org/10.1159/000096090

Pardon, M.-C., Gérardin, P., Joubert, C., Pérez-Diaz, F., Cohen-Salmon, C., 2000. Influence of prepartum chronic ultramild stress on maternal pup care behavior in mice. Biol. Psychiatry 47, 858–863. https://doi.org/10.1016/S0006-3223(99)00253-X

Pawluski, J.L., Lambert, K.G., Kinsley, C.H., 2016. Neuroplasticity in the maternal hippocampus: Relation to cognition and effects of repeated stress. Horm. Behav., Parental Care 77, 86–97. https://doi.org/10.1016/j.yhbeh.2015.06.004

Pawluski, J.L., Lonstein, J.S., Fleming, A.S., 2017. The Neurobiology of Postpartum Anxiety and Depression. Trends Neurosci. 40, 106–120. https://doi.org/10.1016/j.tins.2016.11.009

Pawluski, J.L., van den Hove, D.L.A., Rayen, I., Prickaerts, J., Steinbusch, H.W.M., 2011. Stress and the pregnant female: Impact on hippocampal cell proliferation, but not affective-like behaviors. Horm. Behav. 59, 572–580. https://doi.org/10.1016/j.yhbeh.2011.02.012

Perani, C.V., Slattery, D., 2014. Using animal models to study post-partum psychiatric disorders. Br. J. Pharmacol. https://doi.org/10.1111/bph.12640

Picard, M., McEwen, B.S., Epel, E.S., Sandi, C., 2018. An energetic view of stress: Focus on mitochondria. Front. Neuroendocrinol. 49, 72–85. https://doi.org/10.1016/j.yfrne.2018.01.001

Post, C., Leuner, B., 2019. The maternal reward system in postpartum depression. Arch. Womens Ment. Health 22, 417–429. https://doi.org/10.1007/s00737-018-0926-y

Qiu, W., Hodges, T.E., Clark, E.L., Blankers, S.A., Galea, L.A.M., 2020. Perinatal depression: Heterogeneity of disease and in animal models. Front. Neuroendocrinol. 59, 100854. https://doi.org/10.1016/j.yfrne.2020.100854

Rangaraju, V., Lauterbach, M., Schuman, E.M., 2019. Spatially Stable Mitochondrial Compartments Fuel Local Translation during Plasticity. Cell 176, 73–84.e15. https://doi.org/10.1016/j.cell.2018.12.013

Ray, S., Tzeng, R.-Y., DiCarlo, L.M., Bundy, J.L., Vied, C., Tyson, G., Nowakowski, R., Arbeitman, M.N., 2016. An Examination of Dynamic Gene Expression Changes in the Mouse Brain During Pregnancy and the Postpartum Period. G3 GenesGenomesGenetics 6, 221–233. https://doi.org/10.1534/g3.115.020982

Rettberg, J.R., Yao, J., Brinton, R.D., 2014. Estrogen: A master regulator of bioenergetic systems in the brain and body. Front. Neuroendocrinol. 35, 8–30. https://doi.org/10.1016/j.yfrne.2013.08.001

Rezin, G.T., Cardoso, M.R., Gonçalves, C.L., Scaini, G., Fraga, D.B., Riegel, R.E., Comim, C.M., Quevedo, J., Streck, E.L., 2008. Inhibition of mitochondrial respiratory chain in brain of rats subjected to an experimental model of depression. Neurochem. Int. 53, 395–400. https://doi.org/10.1016/j.neuint.2008.09.012

Riley, J.S., Tait, S.W., 2020. Mitochondrial DNA in inflammation and immunity. EMBO Rep. 21. https://doi.org/10.15252/embr.201949799

Ross, L.E., Evans, S.E.G., Sellers, E.M., Romach, M.K., 2003. Measurement issues in postpartum depression part 1: Anxiety as a feature of postpartum depression. Arch. Womens Ment. Health 6, 51–57. https://doi.org/10.1007/s00737-002-0155-1

Schiller, C.E., Meltzer-Brody, S., Rubinow, D.R., 2015. The Role of Reproductive Hormones in Postpartum Depression. CNS Spectr. 20, 48–59. https://doi.org/10.1017/S1092852914000480

Smith, J.W., Seckl, J.R., Evans, A.T., Costall, B., Smythe, J.W., 2004. Gestational stress induces post-partum depression-like behaviour and alters maternal care in rats. Psychoneuroendocrinology 29, 227–244. https://doi.org/10.1016/s0306-4530(03)00025-8

van der Kooij, M.A., Hollis, F., Lozano, L., Zalachoras, I., Abad, S., Zanoletti, O., Grosse, J., Guillot de Suduiraut, I., Canto, C., Sandi, C., 2018. Diazepam actions in the VTA enhance social dominance and mitochondrial function in the nucleus accumbens by activation of dopamine D1 receptors. Mol. Psychiatry 23, 569–578. https://doi.org/10.1038/mp.2017.135

Walton, N., Maguire, J., 2019. Allopregnanolone-based treatments for postpartum depression: Why/how do they work? Neurobiol. Stress 11, 100198. https://doi.org/10.1016/j.ynstr.2019.100198

Weger, M., Sandi, C., 2018. High anxiety trait: A vulnerable phenotype for stress-induced depression. Neurosci. Biobehav. Rev. 87, 27–37. https://doi.org/10.1016/j.neubiorev.2018.01.012

West, A.P., 2017. Mitochondrial dysfunction as a trigger of innate immune responses and inflammation. Toxicology, Mitochondrial Toxicity 391, 54–63. https://doi.org/10.1016/j.tox.2017.07.016

West, A.P., Shadel, G.S., 2017. Mitochondrial DNA in innate immune responses and inflammatory pathology. Nat. Rev. Immunol. 17, 363–375. https://doi.org/10.1038/nri.2017.21

Willner, P., 2017. The chronic mild stress (CMS) model of depression: History, evaluation and usage. Neurobiol. Stress, SI:Stressors in animals 6, 78–93. https://doi.org/10.1016/j.ynstr.2016.08.002

Willner, P., Towell, A., Sampson, D., Sophokleous, S., Muscat, R., 1987. Reduction of sucrose preference by chronic unpredictable mild stress, and its restoration by a tricyclic antidepressant. Psychopharmacology (Berl.) 93, 358–364. https://doi.org/10.1007/BF00187257

Workman, J.L., Brummelte, S., Galea, L. a. M., 2013. Postpartum Corticosterone Administration Reduces Dendritic Complexity and Increases the Density of Mushroom Spines of Hippocampal CA3 Arbours in Dams. J. Neuroendocrinol. 25, 119–130. https://doi.org/10.1111/j.1365-2826.2012.02380.x

Yang, S.-H., Liu, R., Perez, E.J., Wen, Y., Stevens, S.M., Valencia, T., Brun-Zinkernagel, A.-M., Prokai, L., Will, Y., Dykens, J., Koulen, P., Simpkins, J.W., 2004. Mitochondrial localization of estrogen receptor. Proc. Natl. Acad. Sci. 101, 4130–4135. https://doi.org/10.1073/pnas.0306948101

Yang, S.-H., Sarkar, S.N., Liu, R., Perez, E.J., Wang, X., Wen, Y., Yan, L.-J., Simpkins, J.W., 2009. Estrogen Receptor β as a Mitochondrial Vulnerability Factor. J. Biol. Chem. 284, 9540–9548. https://doi.org/10.1074/jbc.M808246200

Zhang, Y., Zhang, L., Shi, B., Huang, F., Gao, Y., Miao, Z., Ma, K., Zhan, Z., Zou, W., Liu, M., 2022. Comparison of the chronic unpredictable mild stress and the maternal separation in mice postpartum depression modeling. Biochem. Biophys. Res. Commun. 632, 24– 31. https://doi.org/10.1016/j.bbrc.2022.09.063

Zhou, R., Yazdi, A.S., Menu, P., Tschopp, J., 2011. A role for mitochondria in NLRP3 inflammasome activation. Nature 469, 221–225. https://doi.org/10.1038/nature09663

Zoubovsky, S.P., Hoseus, S., Tumukuntala, S., Schulkin, J.O., Williams, M.T., Vorhees, C.V., Muglia, L.J., 2020. Chronic psychosocial stress during pregnancy affects maternal behavior and neuroendocrine function and modulates hypothalamic CRH and nuclear steroid receptor expression. Transl. Psychiatry 10, 6. https://doi.org/10.1038/s41398-020-0704-2

