## Supplementary figures and images for "Gestational stress decreases postpartum mitochondrial respiration in the prefrontal cortex of female rats"

### Supplemental Figure 1

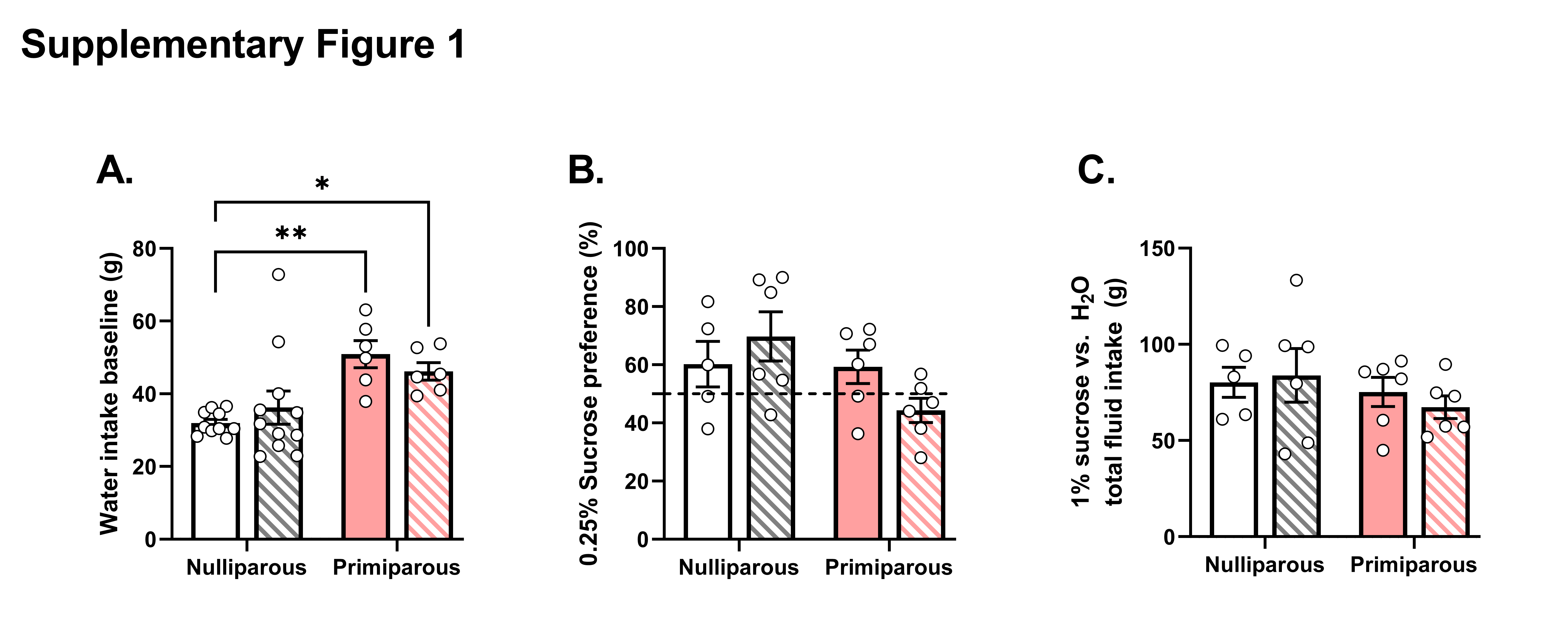

### Supplemental Figure 2

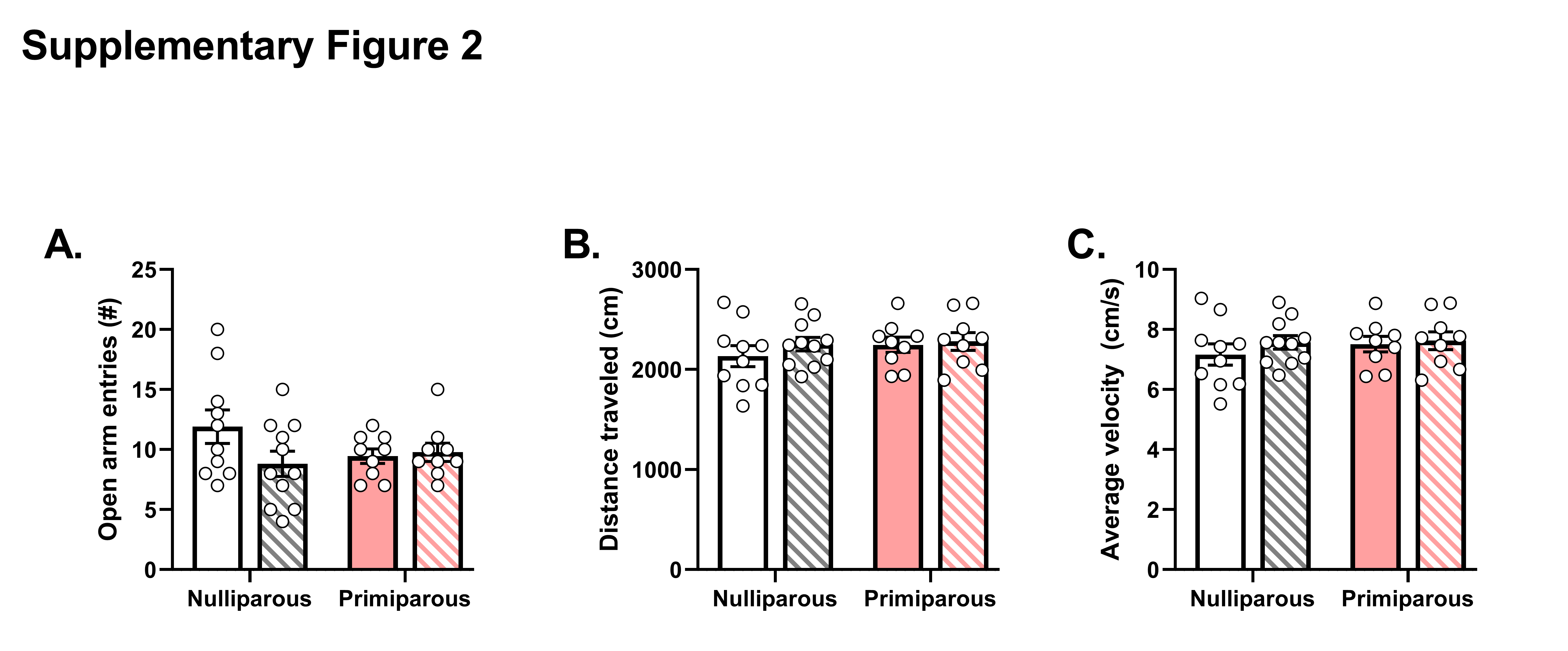

### Supplemental Figure 3

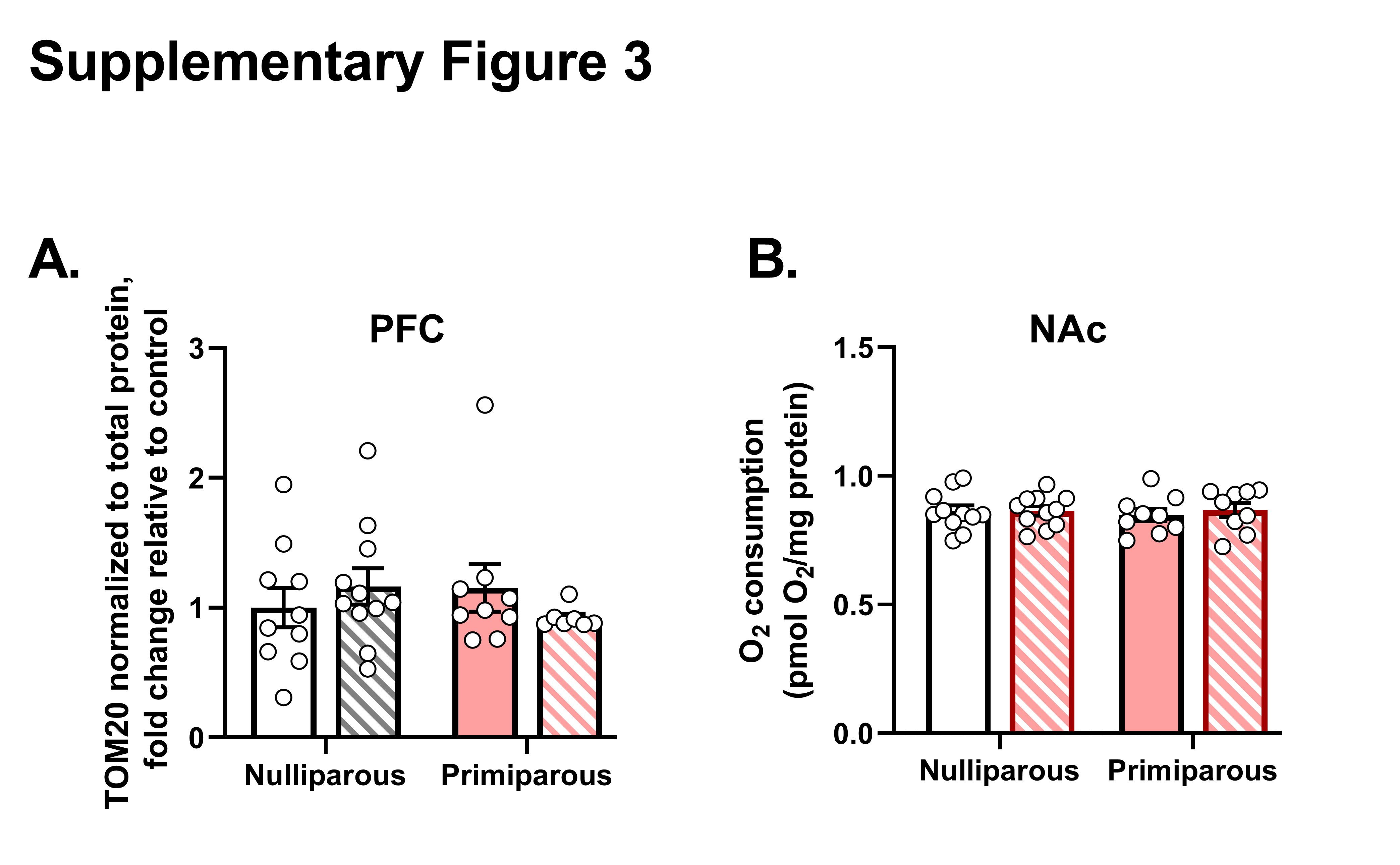

### Supplemental Figure 4

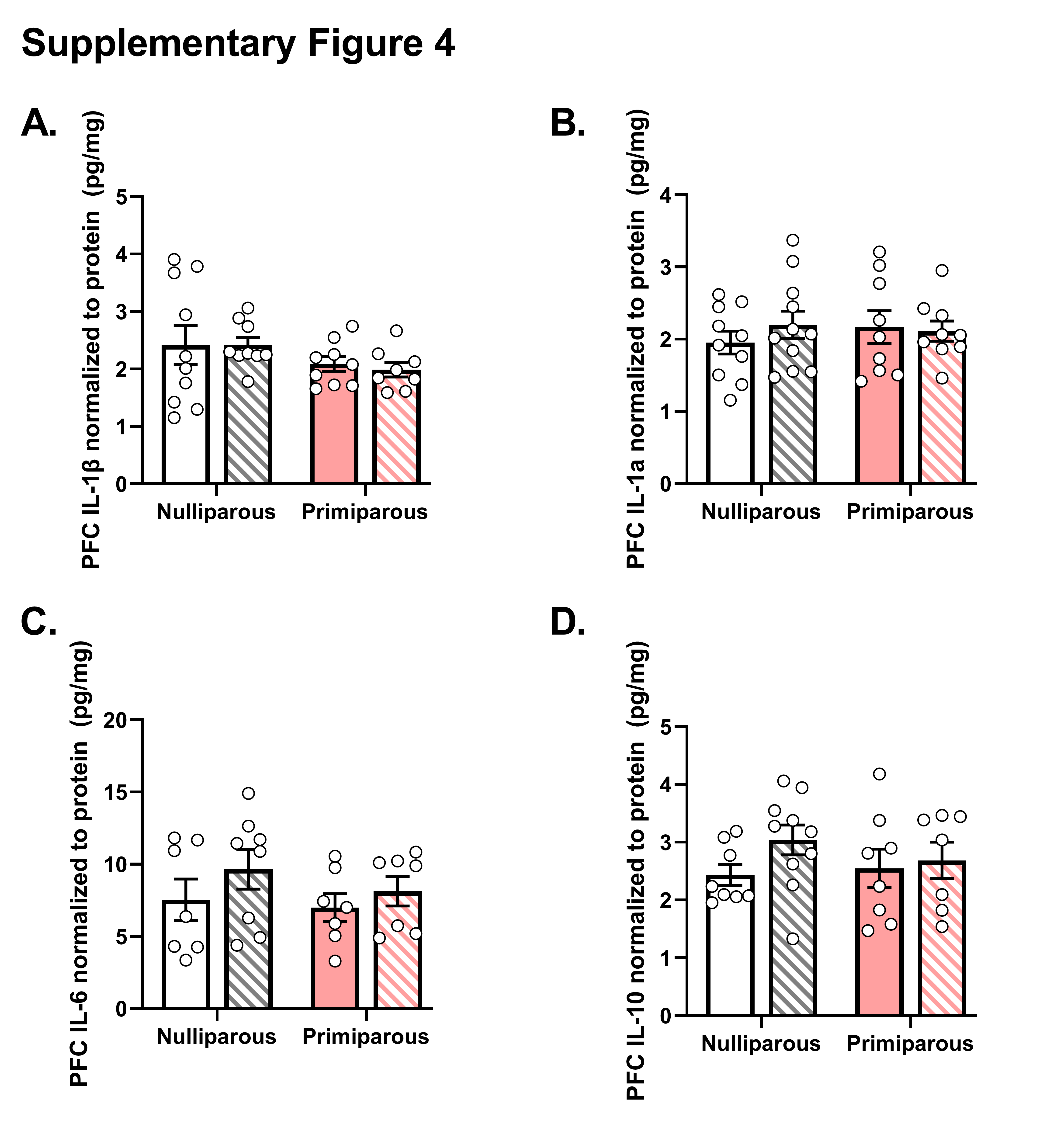

### Supplemental Figure 5

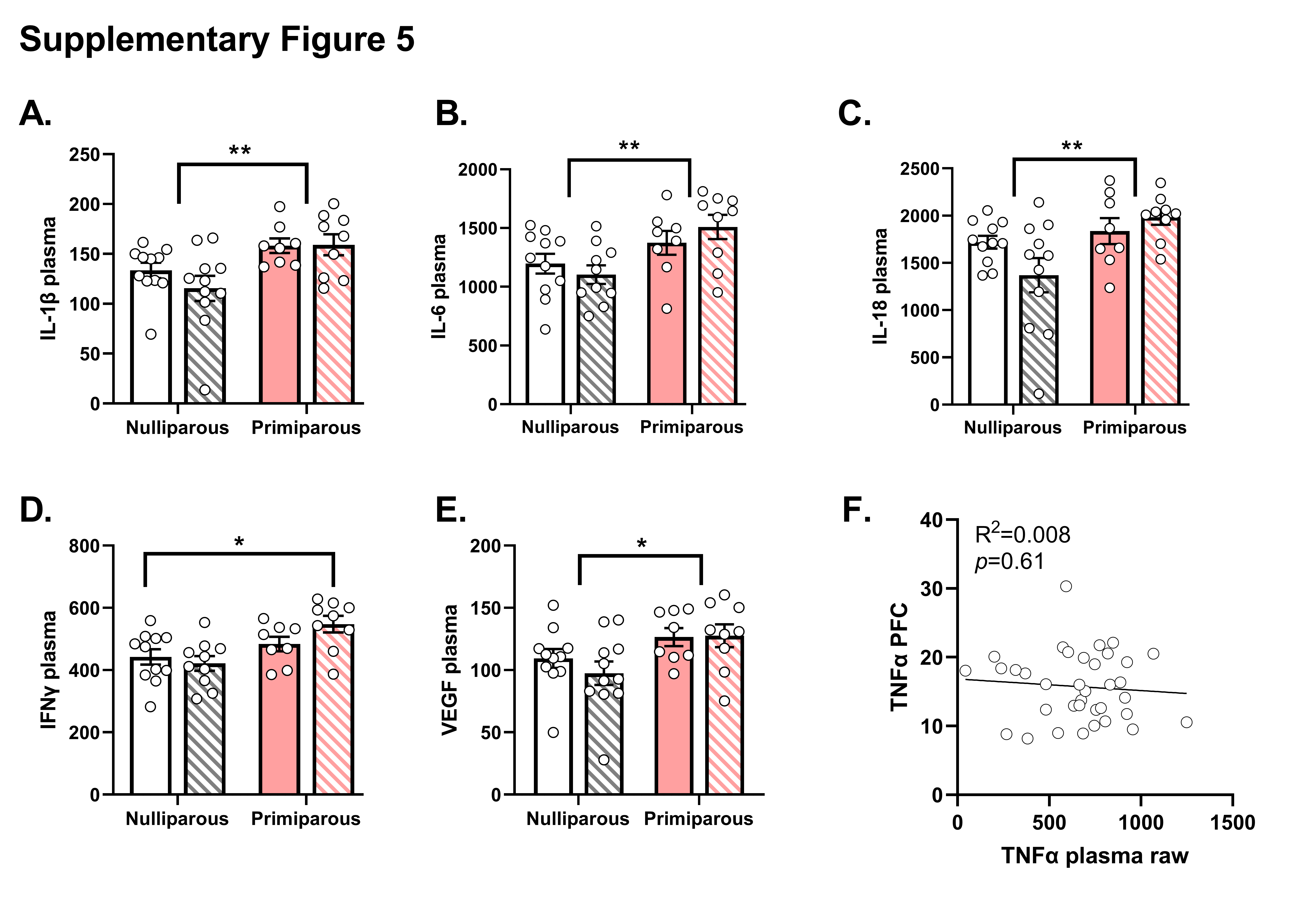
